# Neogenin-1 distinguishes between myeloid-biased and balanced *Hoxb5*^+^ mouse long-term hematopoietic stem cells

**DOI:** 10.1101/608398

**Authors:** Gunsagar S. Gulati, Monika Zukowska, Joseph Noh, Allison Zhang, Rahul Sinha, Benson M. George, Daniel J. Wesche, Irving L. Weissman, Krzysztof Szade

## Abstract

Hematopoietic stem cells (HSCs) self-renew and generate all blood cells. Recent studies with single-cell transplants (1–3) and lineage tracing (4, 5) suggest that adult HSCs are diverse in their reconstitution and lineage potentials. However, prospective isolation of these subpopulations has remained challenging. Here, we identify Neogenin-1 (NEO1) as a unique surface marker on a fraction of mouse HSCs labeled with *Hoxb5*, a specific reporter of long-term HSCs (LT-HSCs) (6). We show that NEO1^+^*Hoxb5*^+^ LT-HSCs expand with age and respond to myeloablative stress, while NEO1^−^*Hoxb5*^+^ LT-HSCs exhibit no significant change in number. NEO1^+^*Hoxb5*^+^ LT-HSCs are more often in the G_2_/S cell cycle phase compared to NEO1^−^*Hoxb5*^+^ LT-HSCs in both young and old bone marrow. Upon serial transplantation, NEO1^+^*Hoxb5*^+^ LT-HSCs exhibit myeloid-biased differentiation and reduced reconstitution, while NEO1^−^*Hoxb5*^+^ LT-HSCs are lineage-balanced and stably reconstitute recipients. Gene expression comparison reveals increased expression of cell cycle genes and evidence of lineage-priming in the NEO1^+^ fraction. Finally, transplanted NEO1^+^*Hoxb5*^+^ LT-HSCs rarely generate NEO1^−^*Hoxb5*^+^ LT-HSCs, while NEO1^−^*Hoxb5*^+^ LT-HSCs repopulate both LT-HSC fractions. This supports a model in which dormant, balanced, NEO1^−^*Hoxb5*^+^ LT-HSCs can hierarchically precede active, myeloid-biased NEO1^+^*Hoxb5*^+^ LT-HSCs.

**SIGNIFICANCE STATEMENT:** Hematopoietic stem cells (HSCs) are rare cells that have the unique ability to regenerate themselves and produce all blood cells throughout life. However, HSCs are functionally heterogeneous and several studies have shown that HSCs can differ in their contribution to major blood lineages. In this study, we discovered that the surface marker, Neogenin-1, can divide mouse HSCs into two subpopulations—one that is more active but biased towards producing myeloid cells and another that is more dormant and capable of equally producing all blood lineages. Neogenin-1 reveals the diversity and hierarchical relationship of HSCs in the mouse bone marrow, enables the prospective isolation of myeloid-biased and balanced HSCs, and opens opportunities to do the same in humans.

## INTRODUCTION

The hematopoietic system is hierarchically organized into distinct cell types and cellular states with unique functions and regenerative potentials (7). Residing at the apex of this hierarchy is the hematopoietic stem cell – the master orchestrator of all blood and immune development, maintenance, and regeneration. HSCs have the unique ability to self-renew and give rise to all major lineages of blood and immune cells throughout life. Over the years, combinations of surface markers (8–12) and reporter genes (6, 13–15) have refined the definition of mouse HSCs and enabled the purification of long-term hematopoietic stem cells (LT-HSCs), a refined subset of HSCs capable of serially reconstituting irradiated recipients in a transplantation model. Recently, we identified *Hoxb5* as a specific marker of long-term repopulating cells and generated a *Hoxb5-*mCherry reporter mouse strain for the prospective isolation of these cells (6). We demonstrated that the *Hoxb5*-mCherry reporter significantly improves the precise isolation of serially transplantable LT-HSCs compared to immunophenotypically defined HSCs (phenotypic HSCs, or pHSCs). As previously shown, only 7-35% of pHSCs are *Hoxb5-*mCherry^+^ and the potential for long-term multilineage reconstitution is restricted to this fraction (6).

Although *Hoxb5*^+^ LT-HSCs are multilineage contributing, self-renewing cells (6), the functional heterogeneity within this compartment has not yet been characterized. Understanding the composition of LT-HSCs may offer valuable insights into the mechanism of HSC expansion with age, as well the competition of diverse HSCs for bone marrow niches (16, 17). On a per cell basis, HSCs from older mice exhibit biased differentiation towards myeloid lineages and reduced stem cell activity, presumably due to weaker responses to SDF-1 (17, 18) and poorer engraftability of HSCs in G_1_ (19) and S/G_2_/M (20). These aged HSCs may arise from either the cell-intrinsic transition of balanced to myeloid-biased LT-HSCs or the clonal expansion of pre-existing fractions of myeloid-biased LT-HSCs (16, 17, 21–23). Several studies support the presence of pre-existing myeloid-biased LT-HSCs by demonstrating that myeloid-biased subpopulations of LT-HSCs in young, healthy mice respond to environmental challenges, such as inflammation and infection (24, 25). Results from lineage tracing with genetic barcodes (4, 5) and single cell transplants of LT-HSCs also support the notion of inherent functional diversity among long-term repopulating HSCs (1–3). However, these studies did not identify markers to prospectively isolate distinct sub-populations of HSCs. Other groups have identified markers, such as CD150 (21), CD41 (26), vWF (27), and CD61 (24) that enrich for self-renewing lineage-biased subpopulations of HSCs. However, these markers were shown to segregate fractions of pHSCs, which contain both short-term, *Hoxb5*^−^ and long-term, *Hoxb5*^+^ HSCs. Our previous study showed that *Hoxb5*^−^ pHSCs are homogenously lymphoid-biased (6), but the diversity of *Hoxb5*^+^ LT-HSCs has not yet been fully explored. Therefore, we sought to interrogate the heterogeneity among purified *Hoxb5*^+^ LT-HSCs and identify a strategy to prospectively isolate these cells with phenotypic markers.

Here, we find that Neogenin-1 (NEO1), a transmembrane receptor of the immunoglobulin family (28), is expressed on a fraction of *Hoxb5*^+^ LT-HSCs and decreases with differentiation. Although NEO1 has been extensively investigated as a receptor for axon guidance (29, 30), neuronal survival (31), skeletal myofiber differentiation (32), intracellular iron homeostasis (33), mammary epithelial development (34), and endothelial migration (35), as of yet, little is known about the cells that express this marker in the bone marrow. We find that NEO1^+^*Hoxb5*^+^ LT-HSCs represent a myeloid-biased subset of LT-HSCs that responds to myeloablative stress and expands with age. Contrastingly, NEO1^−^*Hoxb5*^+^ LT-HSCs exhibit greater reconstitution potential, balanced lineage output, and a more quiescent cell cycle status compared to NEO1^+^*Hoxb5*^+^ LT-HSCs. After transplant, NEO1^−^*Hoxb5*^+^ LT-HSCs give rise to NEO1^+^*Hoxb5*^+^ LT-HSCs, but the reverse transition is rarely observed. We, therefore, propose a model of early long-term hematopoiesis in which balanced, quiescent LT-HSCs self-renew and generate long-term myeloid-biased LT-HSCs in response to stress and during the course of aging.

## RESULTS

### Neogenin-1 (NEO1) marks a subpopulation of mouse *Hoxb5*^+^ LT-HSCs and human HSCs

Functional heterogeneity within *Hoxb5*^+^ LT-HSCs is poorly understood. To identify surface candidates that fractionate *Hoxb5*^+^ LT-HSCs, we first pattern-searched 64 microarray expression profiles of 23 distinct mouse hematopoietic cell types (36) for (1) genes annotated to code for cell surface proteins (GO Biological Process: 0009986) and (2) genes specifically expressed in HSCs compared to downstream cell types (Fig. 1*A*; *SI Appendix*, Table S1). We found several known HSC-specific markers, including *Robo4* (37), *Slamf1* (11), *Ly6a* (8)*, Vwf* (27)*, TEK* (38), and a member of the *Gpcr5* family (39), validating the utility of our approach. We also identified several novel markers of HSCs that have not been previously reported (Fig. 1*A*). Among the top 3 most enriched surface markers on HSCs, Neogenin-1 (*Neo1*) was more highly expressed on HSCs compared to the other two candidates (Fig. 1 *A* and *B*). Single-cell RNA-sequencing data of hematopoietic stem and progenitor cells validated the enriched expression of *Neo1* in LT-HSCs compared to downstream short-term HSCs and progenitors (*SI Appendix,* Fig. S1). *Neo1* was also expressed on subsets of bone marrow stromal and endothelial cells and demarcated the contours of trabecular bone (*SI Appendix,* Fig. S2), suggesting its expression is not restricted to hematopoietic cells in the bone marrow.

**Figure 1.**
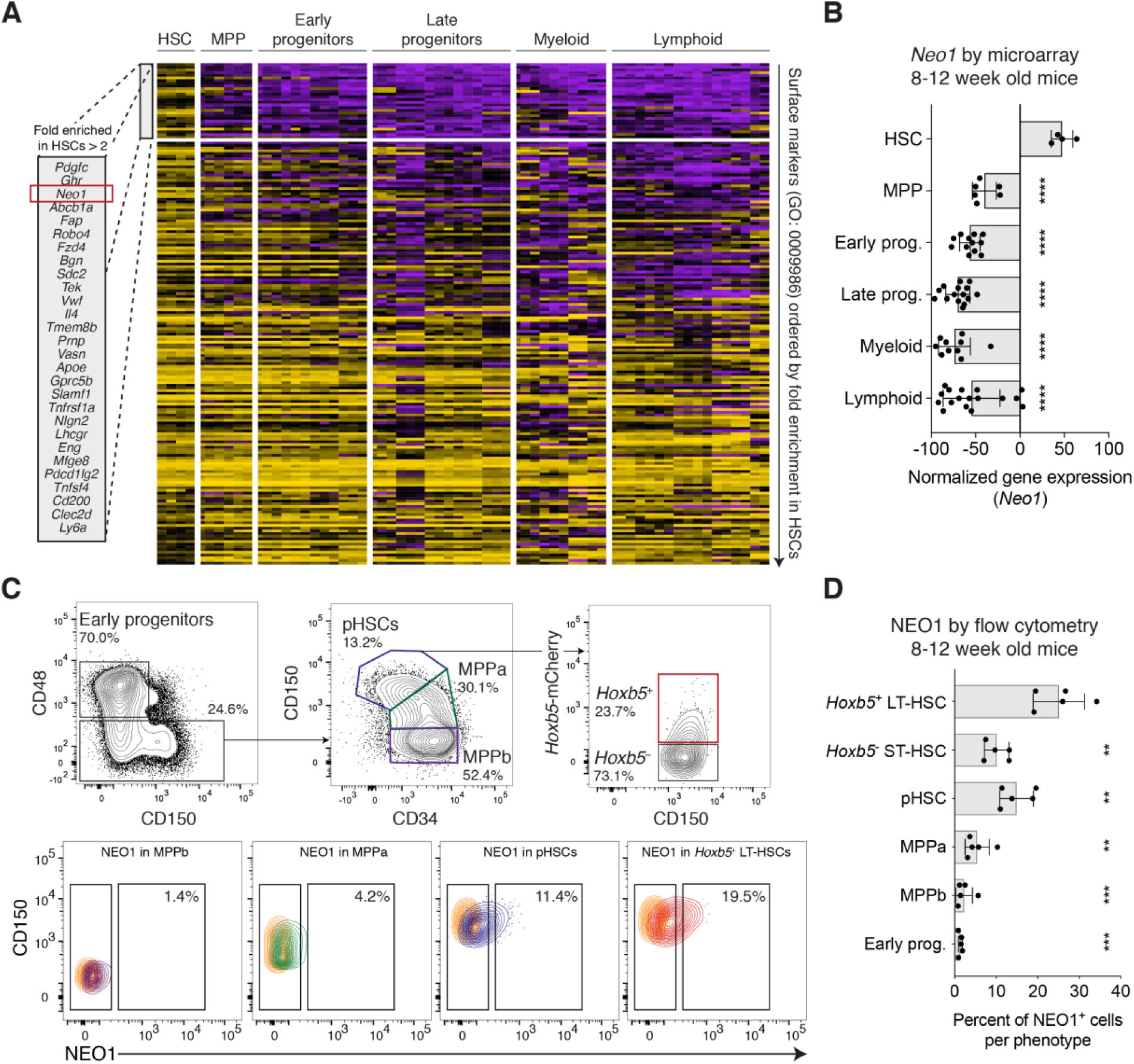
Identification of Neogenin-1 as a unique surface marker on mouse hematopoietic stem cells (HSCs). (**A and B**) *In silico* screen to identify unique surface markers on HSCs from 64 microarray expression profiles of 23 distinct mouse hematopoietic cell types. (**A**) Heatmap showing normalized gene expression of gene-ontology-annotated (GO: 0009986) surface markers across different hematopoietic compartments. 186 surface markers expressed on HSCs and 23 phenotypes categorized as ‘HSCs’, ‘MPPs’, ‘Early progenitors’, ‘Late progenitors’, ‘Myeloid’, and ‘Lymphoid’ are displayed (*SI Appendix*, Table S1). Genes are ordered from top to bottom by log_2_ fold enrichment in HSCs compared to downstream cells and the top most enriched genes (>2 log_2_ fold enrichment) are highlighted in a box. For further details, see methods. (**B**) Barplots showing normalized gene expression of *Neo1* across the cell type categories shown in **A**. Statistical significance was calculated by unpaired, two-tailed Student’s *t*-test between ‘HSC’ and each cell type. \*\*\*\**P* < 0.0001. (**C and D**) Flow cytometry analysis of NEO1 surface expression in the mouse bone marrow (*n* = 5 mice). (**C**) Contour plots with outliers showing the gating scheme for early progenitors, MPPb, MPPa, pHSCs, *Hoxb5*^−^ ST-HSCs, and *Hoxb5*^+^ LT-HSCs (top) and the corresponding surface expression of NEO1 for select populations (bottom). Colors correspond to populations shown. Goat IgG isotype control for fluorescence staining with goat anti-mouse/human NEO1 antibody is highlighted in orange (bottom). (**D**) Barplots showing the percent of NEO1^+^ cells for each cell type gated in **c**. Statistical significance was calculated by a paired, two-tailed Student’s *t*-test between ‘*Hoxb5*^+^ LT-HSC’ and each cell type. \*\**P* < 0.01, \*\*\**P* < 0.001. Barplots in this figure indicate mean ± SD.

We next used flow cytometry to measure the relative protein levels of NEO1 on the surface of 2- to-3-month-old early hematopoietic progenitors, multipotent progenitor subset A (MPPa), multipotent progenitor subset B (MPPb), phenotypic HSCs defined as Lin^−^c-KIT^+^SCA1^+^CD48^−^ FLK2^−^CD150^+^CD34^−^ (hereafter referred to as pHSCs), and two populations among pHSCs, including *Hoxb5*^+^ LT-HSCs and *Hoxb5*^−^ short-term HSCs (ST-HSCs) (Fig. 1*C*; *SI Appendix,* Fig. S3). Consistent with its gene expression, the relative protein levels of NEO1 and the frequency of NEO1^+^ cells progressively decreased with differentiation. NEO1^+^ cells comprised a significantly higher fraction of *Hoxb5*^+^ LT-HSCs compared to downstream cells (Fig. 1 *C* and *D*). NEO1 was also expressed on a fraction of long-term reconstituting Lin^−^CD34^+^CD38^−^ CD45RA^−^CD90^+^ HSCs from human bone marrow (40), although NEO1 enrichment in human HSCs was diminished compared to that observed in mouse HSCs (*SI Appendix*, Fig. S4).

### NEO1^+^*Hoxb5*^+^ LT-HSCs selectively expand with age and respond to myeloablative stress

Previous studies have shown that a subpopulation of pHSCs expands with age (2, 21) and responds to environmental challenge (24, 25). However, the effect of aging and stress on LT-HSCs and their subpopulations have not yet been evaluated. To that end, we first measured the number and frequency of *Hoxb5*^+^ LT-HSCs (Fig. 2 *A* to *C*; *SI Appendix,* Fig. S5) and NEO1^+^ and NEO1^−^ fractions (Fig. 2 *D* to *F*; *SI Appendix,* Fig. S5) in 2-month, 5-month, 13-month, and 22-month-old bone marrow. Consistent with the overall expansion of pHSCs (*SI Appendix,* Fig. S5 *A* and *B*), we observed that the total number of *Hoxb5*^+^ LT-HSCs and *Hoxb5*^−^ ST-HSCs was significantly increased (Fig. 2 *B* and *C*; *SI Appendix,* Fig. S5 *C* to *E*). The frequency of *Hoxb5*^+^ LT-HSCs among pHSCs, although on average higher in bone marrow from older (13-month-old and 22-month-old) than younger (2-month-old and 5-month-old) mice, was highly variable in aged mice (Fig. 2*A*; *SI Appendix,* Fig. S5*C*).

**Figure 2.**
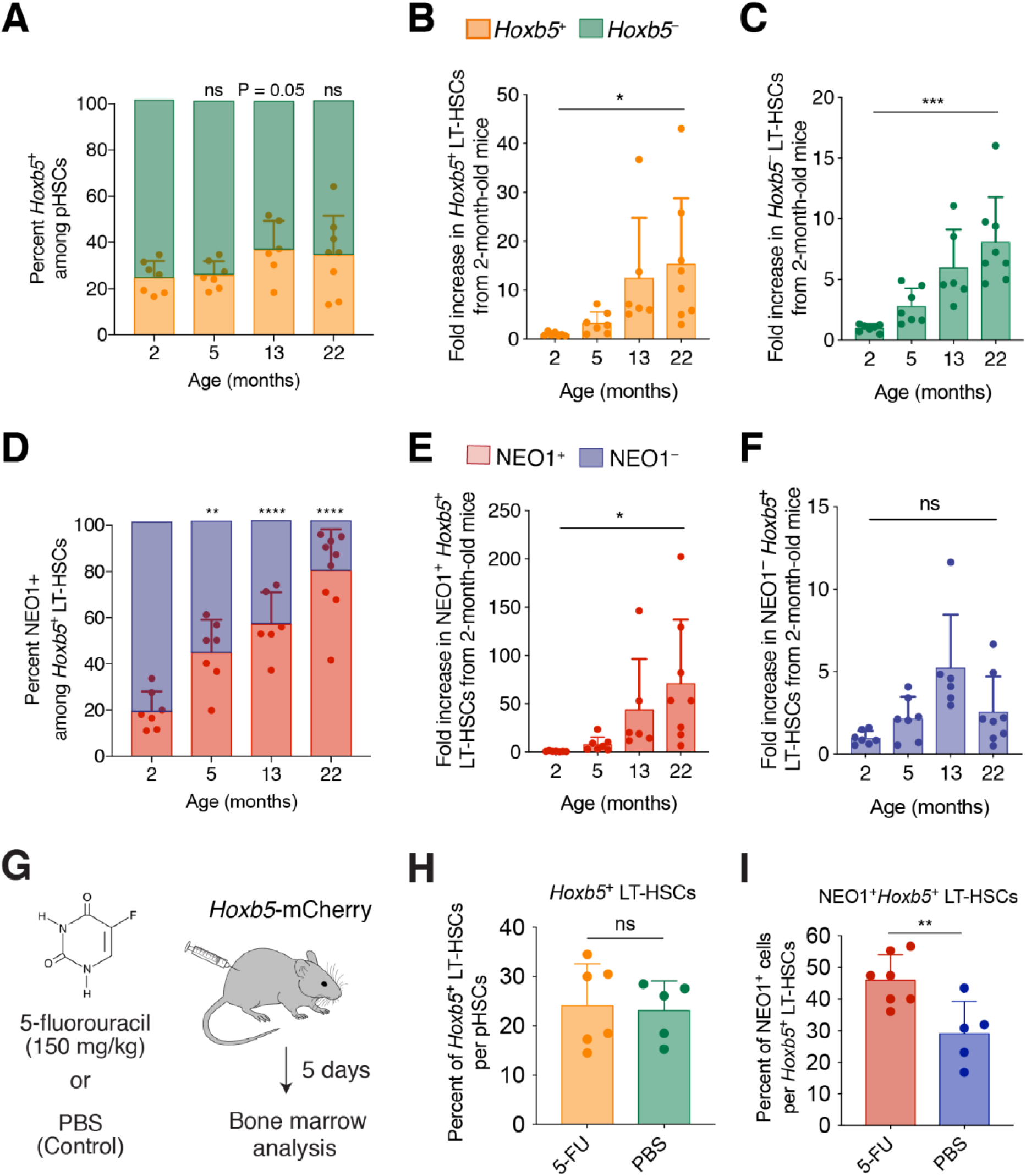
NEO1^+^*Hoxb5*^+^ LT-HSCs selectively expand during aging and respond to myeloablative stress. (**A to C**) Frequency and number of *Hoxb5*^+^ LT-HSCs and *Hoxb5*^−^ ST-HSCs at 2 (*n* = 7 mice), 5 (*n* = 7 mice), 13 (*n* = 6 mice), and 22 (*n* = 9 mice) months of age. (**A**) Percent of cells among pHSCs that are *Hoxb5*^+^ (orange) and Hoxb5^−^ (green). Statistical significance was calculated using an unpaired, two-tailed Student’s *t*-test between 2 months and each time point. ns = non-significant, *P* ≥ 0.05. (**B**) Number of *Hoxb5*^+^ LT-HSCs per million whole bone marrow (WBM) cells. Statistical significance was calculated using an unpaired, two-tailed Student’s *t*-test between 2 months and 22 months of age. \**P* < 0.05. (**C**) Number of *Hoxb5*^−^ ST-HSCs per million whole bone marrow (WBM) cells. Statistical significance was calculated using an unpaired, two-tailed Student’s *t*-test between 2 months and 22 months of age. \*\*\**P* < 0.001. (**D to F**) Frequency and number of NEO1^+^ and NEO1^−^*Hoxb5*^+^ LT-HSCs at 2 (*n* = 7 mice), 5 (*n* = 7 mice), 13 (*n* = 6 mice), and 22 (*n* = 9 mice) months of age. (**D**) Percent of cells among *Hoxb5*^+^ LT-HSCs that are NEO1^+^ (red) and NEO1^−^ (blue). Statistical significance was calculated using an unpaired, two-tailed Student’s *t*-test between 2 months and each time point. \*\*\**P* < 0.001, \*\*\*\**P* < 0.0001 (**E**) Number of NEO1^+^ *Hoxb5*^+^ LT-HSCs per million whole bone marrow (WBM) cells. Statistical significance was calculated using an unpaired, two-tailed Student’s *t*-test between 2 months and 22 months of age. \**P* < 0.05. (**F**) Number of NEO1^−^ *Hoxb5*^+^ LT-HSCs per million whole bone marrow (WBM) cells. Statistical significance was calculated using an unpaired, two-tailed Student’s *t*-test between 2 months and 22 months of age. ns = non-significant, *P* ≥ 0.05. (**G to I**) Response of HSC subpopulations from 4-month-old *Hoxb5-*mCherry mice 5 days after treatment with 5-fluoruracil (5-FU). (**G**) Experimental design of myeloablative stress with 5-FU (*n* = 6 mice) with PBS control (*n* = 5 mice). (**H**) Frequency of *Hoxb5*^+^ LT-HSCs and *Hoxb5*^−^ ST-HSCs among all pHSCs 5 days after treatment with 5-FU or PBS. Statistical significance was calculated using an unpaired, two-tailed Student’s *t*-test. ns = non-significant, *P* ≥ 0.05. (**I**) Frequency of NEO1^+^ and NEO1^−^ *Hoxb5*^+^ LT-HSCs among all *Hoxb5*^+^ LT-HSCs 5 days after treatment with 5-FU or PBS. Statistical significance was calculated using an unpaired, two-tailed Student’s *t*-test. \*\**P* < 0.01. All barplots in this figure indicate mean ± SD.

Despite the variable expansion of *Hoxb5*^+^ LT-HSCs, the frequency of NEO1^+^ cells among *Hoxb5*^+^ LT-HSCs progressively increased with age in a consistent manner (Fig. 2*D*; *SI Appendix,* Fig. S5*F*). Fewer than 20% of *Hoxb5*^+^ LT-HSCs in 2-month-old mice expressed surface NEO1, while more than 80% of 22-month-old *Hoxb5*^+^ LT-HSCs were NEO1^+^ (Fig. 2*D*; *SI Appendix,* Fig. S5*F*). However, while the number of NEO1^+^ cells per million whole bone marrow cells increased with age (Fig. 2*E*; *SI Appendix,* Fig. S5*G*), the number of NEO1^−^*Hoxb5*^+^ LT-HSCs did not significantly change (Fig. 2*F*; *SI Appendix,* Fig. S5*H*). This suggests that the NEO1^+^ fraction selectively expands among *Hoxb5*^+^ LT-HSCs in the bone marrow, while the number of NEO1^−^ *Hoxb5*^+^ LT-HSCs remains stable with age.

We next evaluated the response of *Hoxb5*^+^ LT-HSCs and the NEO1^+^ and NEO1^−^ subpopulations to myeloablative stress. 4-month-old adult mice were treated with 150 mg/kg 5-fluorouracil (5-FU) and their bone marrow was analyzed 5 days post-treatment when HSC proliferation is maximum. (41) (Fig. 2*G*). While the frequency of *Hoxb5*^+^ and *Hoxb5*^−^ cells among pHSCs did not change (Fig. 2*H*), a significantly higher percentage of *Hoxb5*^+^ LT-HSCs were NEO1^+^ than NEO1^−^ after treatment (Fig. 2*I*). This response may be due to the proliferation of NEO1^+^*Hoxb5*^+^ LT-HSCs, differentiation of NEO1^−^ *Hoxb5*^+^ to NEO1^+^*Hoxb5*^+^, or a combination of the two.

### NEO1^+^*Hoxb5*^+^ LT-HSCs are more proliferative than NEO1^−^*Hoxb5*^+^ LT-HSCs in young and old mice

We next asked whether the difference in expansion of NEO1^+^ versus NEO1^−^*Hoxb5*^+^ LT-HSCs during aging and in response to myeloablative stress can be partially explained by differences in proliferation (42). To address this, we measured the percent of each cell population in G_0_, G_1_, and G_2_/S by KI-67 and DAPI staining at both 2-to-3 months and 12-to-14 months of age (Fig. 3; *SI Appendix,* Fig. S6). Consistent with previous reports (43), 12-to-14-month-old HSCs were more often in G_0_ compared to 2-to-3-month-old HSCs, suggesting increased exhaustion and senescence with age in the absence of leukemia (Fig. 3; *SI Appendix,* Fig. S6). In support of the limited expansion of *Hoxb5*^+^ LT-HSCs among pHSCs with age, we did not observe significant differences in G_0_, G_1_, or G_2_/S status between *Hoxb5*^+^ LT-HSCs and *Hoxb5*^−^ ST-HSCs (Fig. 3 *A* to *C*; *SI Appendix,* Fig. S6). NEO1^+^ and NEO1^−^*Hoxb5*^+^ LT-HSCs also did not differ in the proportion of G_0_ cells in both 2-to-3-month-old and 12-to-14-month-old bone marrow (Fig. 3*D*; *SI Appendix,* Fig. S6)

**Figure 3.**
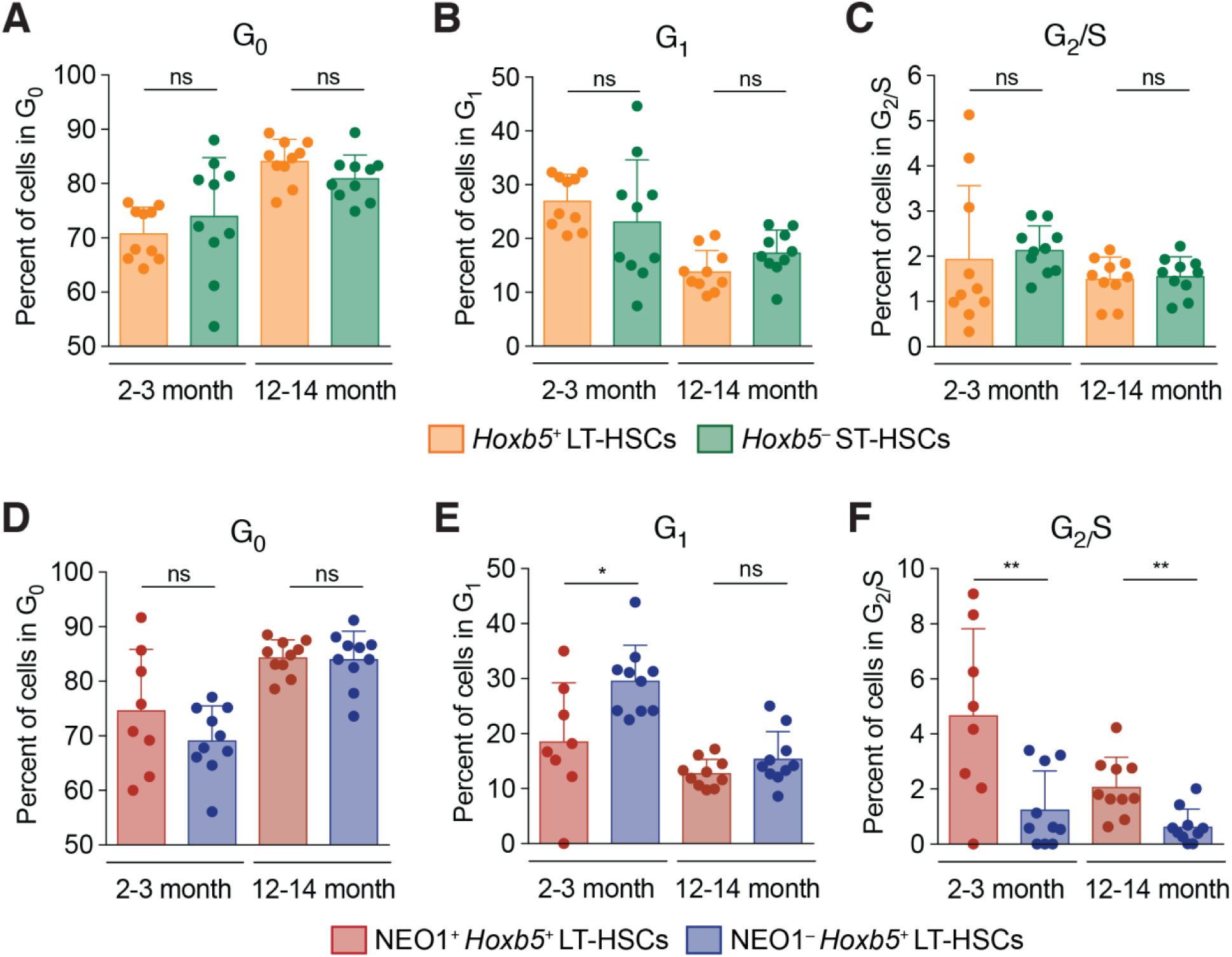
Neogenin-1 marks a more proliferative fraction of LT-HSCs. (**A to F**) Cell cycle analysis of 2-to-3-month-old (*n* = 10 mice) and 12-to-14-month-old (*n* = 10 mice) LT-HSC fractions with Ki-67 and DAPI staining. (**A to C**) Percent of *Hoxb5*^+^ LT-HSCs or *Hoxb5*^−^ ST-HSCs in (**A**) G_0_, (**B**) G_1_, and (**C**) G_2_/S in 2-to-3-month-old and 12-to-14-month-old mouse bone marrow. Statistical significance was calculated using an unpaired, two-tailed Student’s *t*-test. ns = non-significant, *P* ≥ 0.05. (**D to F**) Percent of NEO1^+^ or NEO1^−^ *Hoxb5*^+^ LT-HSCs in (**D**) G_0_, (**E**) G_1_, and (**F**) G_2_/S in 2-to-3-month-old and 12-to-14-month-old mouse bone marrow. Statistical significance was calculated using an unpaired, two-tailed Student’s *t*-test. ns = non-significant, *P* ≥ 0.05, \**P* < 0.05, \*\**P* < 0.01. All barplots in this figure indicate mean ± SD.

However, NEO1^+^*Hoxb5*^+^ LT-HSCs were significantly more often in G_2_/S compared to NEO1^−^ *Hoxb5*^+^ LT-HSCs in both young and old bone marrow (Fig. 3*F*; *SI Appendix,* Fig. S6). Moreover, in the young bone marrow, there was a significantly smaller percentage of NEO1^+^*Hoxb5*^+^ LT-HSCs in G_1_ compared to NEO1^−^*Hoxb5*^+^ LT-HSCs (Fig. 3*E*; *SI Appendix,* Fig. S6). Taken together, this suggests that NEO1^+^*Hoxb5*^+^ LT-HSCs are more proliferative, which may partially contribute to their selective expansion during aging and in response to myeloablative stress.

### Neogenin-1 marks a less regenerative, myeloid-biased fraction of *Hoxb5*^+^ LT-HSCs

Given the aging and cell cycle differences between NEO1^+^ and NEO1^−^*Hoxb5*^+^ LT-HSCs, we next evaluated their reconstitution potential and lineage output by 10-cell transplants into congenic irradiated primary recipients (Fig. 4*A*). Over the course of 16 weeks, the percent of total chimerism among peripheral blood that was donor-derived, was similar between NEO1^+^ and NEO1^−^*Hoxb5*^+^ LT-HSC transplants (Fig. 4*B*). However, among all donor-derived peripheral blood, NEO1^+^*Hoxb5*^+^ LT-HSCs gave rise to a higher percentage of granulocytes and monocytes (myeloid) and lower percentage of B and T cells (lymphoid) compared to NEO1^−^*Hoxb5*^+^ LT-HSCs (Fig. 4 *C* and *D*).

**Figure 4.**
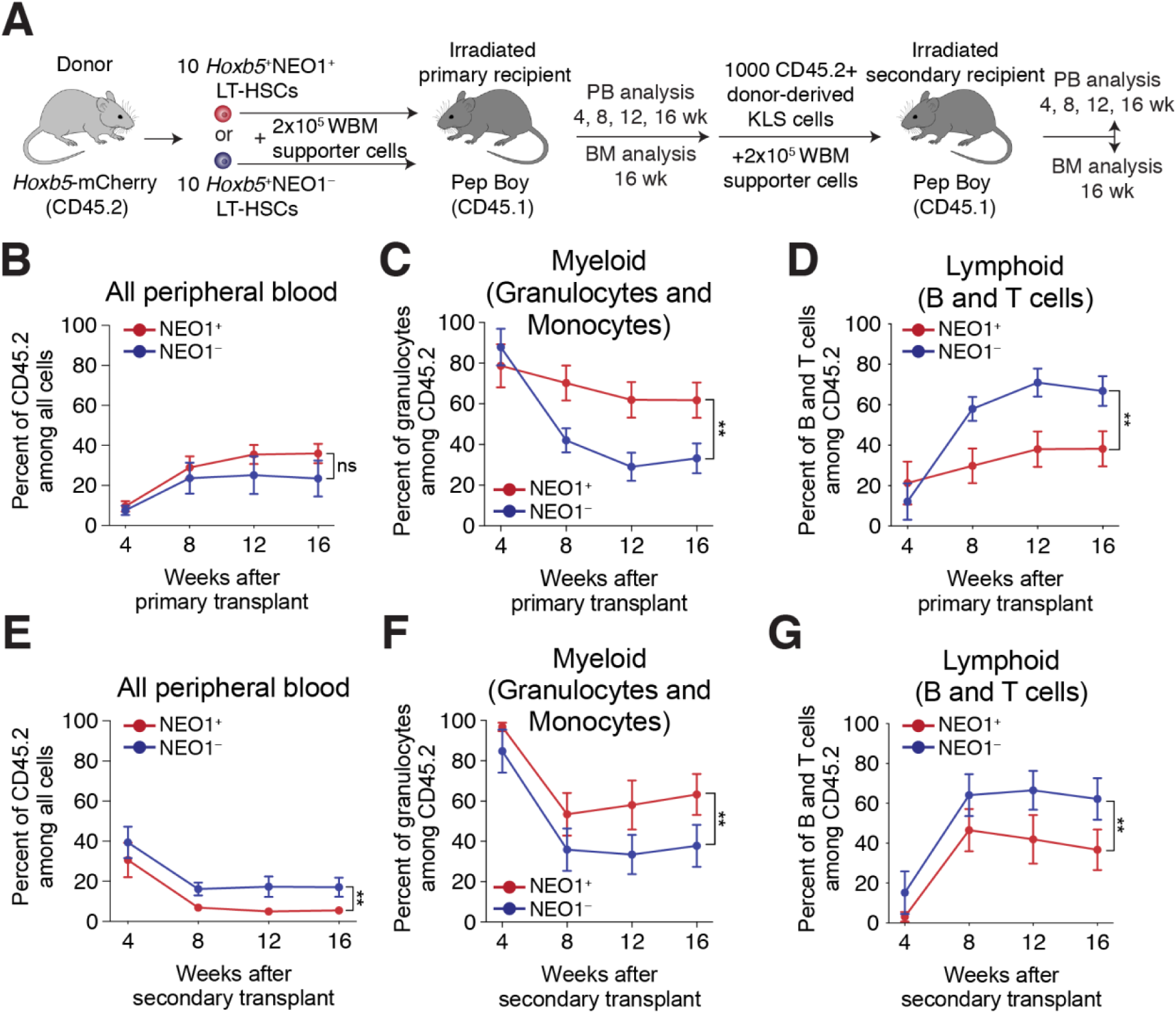
NEO1^+^ *Hoxb5*^+^ LT-HSCs exhibit myeloid bias and reduced reconstitution potential upon serial transplantation. (**A**) Experimental design for primary and secondary transplantations of NEO1^+^ and NEO1^−^*Hoxb5*^+^ LT-HSCs. (**B to D**) Measurements of reconstitution potential and lineage priming after primary transplantation (‘NEO1^+^*’*, *n* = 14 mice; ‘NEO1^−^*’*, *n* = 11 mice). (**B**) Percent of donor-derived (CD45.2^+^) cells among all peripheral blood cells at 4, 8, 12, and 16 weeks post-transplant. (**C**) Percent of myeloid cells (GR1^+^CD11B^+^ granulocytes and monocytes) among donor-derived (CD45.2^+^) cells at 4, 8, 12, and 16 weeks post-transplant. (**D**) Percent of lymphoid cells (B220^+^ B cells and CD3^+^ T cells) among donor-derived (CD45.2^+^) cells at 4, 8, 12, and 16 weeks post-transplant. (**E to G**) Same as in **B to D** but analyzing peripheral blood in secondary recipients transplanted with 1000 donor-derived Lin^−^c-KIT^+^SCA1^+^ (KLS) cells from primary hosts (‘NEO1^+^*’*, *n* = 8; ‘NEO1^−^*’*, *n* = 9). Statistical significance for **B to G** was calculated using two-way ANOVA with time post-transplant and NEO1 status as factors. \*\**P* < 0.01, ns = non-significant. All line plots in this figure indicate mean ± SEM.

To evaluate the long-term reconstitution potential and the stability of lineage bias, we serially transplanted 1000 donor-derived Lin^−^c-KIT^+^SCA1^+^ (KLS) cells from primary recipients into congenic irradiated secondary hosts (Fig. 4*A*). Although KLS cells from both donors repopulated all major lineages in secondary hosts, NEO1^−^*Hoxb5*^+^-derived cells exhibited significantly higher reconstitution compared to NEO1^+^*Hoxb5*^+^-derived cells (Fig. 4*E*). Moreover, as during primary transplant, NEO1^+^*Hoxb5*^+^-derived cells maintained significant bias towards granulocytes and monocytes and away from B and T cells compared to NEO1^−^*Hoxb5*^+^-derived cells. (Fig. 4 *F* and *G*). This suggests that both myeloid-biased and balanced phenotypes are long-term maintained on secondary transplantation.

### Transcriptional programs recapitulate functional differences between NEO1^+^ and NEO1^−^ *Hoxb5*^+^ LT-HSCs

We next sought to understand the transcriptional programs that drive the observed functional differences between NEO1^+^*Hoxb5*^+^ and NEO1^−^*Hoxb5*^+^ LT-HSCs. We isolated 250-500 NEO1^+^*Hoxb5*^+^ and NEO1^−^*Hoxb5*^+^ cells from female, 8-to-12-week old *Hoxb5-*mCherry mice and performed low-input full-length RNA-sequencing using the Smart-Seq2 protocol (44) (Fig. 5*A*). Paired gene expression comparison of the two populations identified 1,036 differentially expressed genes (false discovery rate *P*-*adjusted* < 0.1; Fig. 5*B*; *SI Appendix,* Fig. S7A and Table S2) (45). Genes implicated in activation, cell cycle, and differentiation, such as *Fanca, Fancb*, and *Mycn* (46, 47) were enriched in NEO1^+^*Hoxb5*^+^ LT-HSCs, while genes involved in anti-redox, e.g. *Sod1* (48), and regulation of stem cell potency, e.g. *Klf4* and *Malat1* (49, 50) were enriched in NEO1^−^*Hoxb5*^+^ LT-HSCs (Fig. 5*B*). Gene set enrichment analysis (GSEA) (51) and hypergeometric test with gene ontology (GO): biological processes (52, 53) revealed that the driving differences (FDR < 0.05, P value < 0.05) between NEO1^+^ and NEO1^−^ cells were cell cycle and ribosomal RNA expression (Fig. 5*C*; *SI Appendix,* Fig. S7).

**Figure 5.**
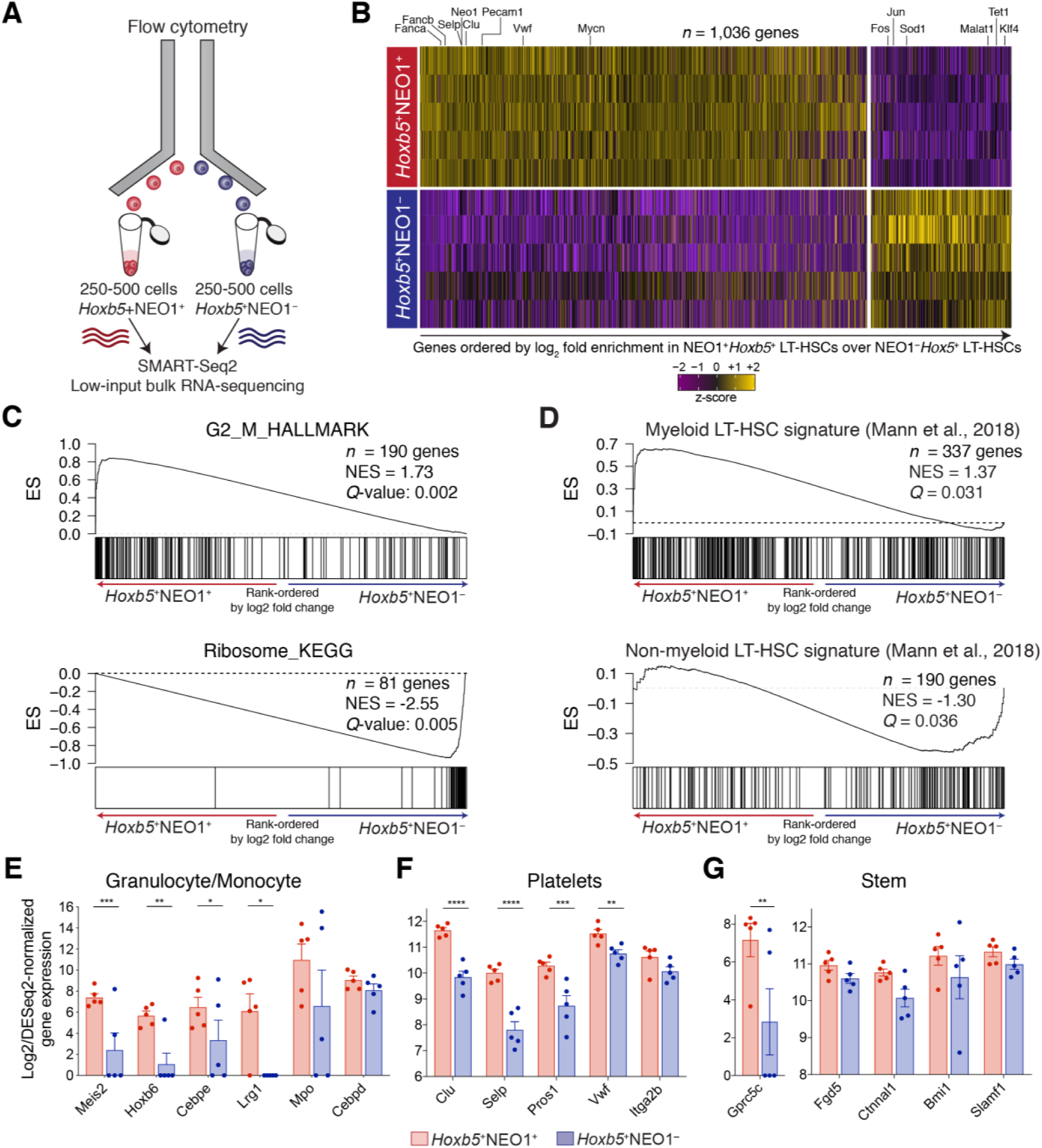
Distinct transcriptional signatures of NEO1^+^ and NEO1^−^ *Hoxb5*^+^LT-HSCs. (**A**) Experimental design for bulk RNA-sequencing of NEO1^+^ and NEO1^−^ *Hoxb5*^+^ LT-HSCs. (**B**) Heatmap of differentially expressed genes (*n =* 1,036 genes; FDR < 0.1) after pairwise comparison of NEO1^+^ (*n* = 5 samples) and NEO1^−^ (*n* = 5 samples) *Hoxb5*^+^ LT-HSC transcriptomes using DESeq2. Select genes are highlighted. Genes are ordered from left to right by log_2_ fold enrichment in NEO1^+^ over NEO1^−^ *Hoxb5*^+^ LT-HSCs. (**C and D**) Gene set enrichment analysis (GSEA) plots of molecular signatures significantly enriched (*Q* value < 0.05) over a gene list ordered by log_2_ fold change, including (**C**) ‘G2_M_HALLMARK’ (*top*), ‘RIBOSOME_KEGG’ (*bottom*), (**D**) Myeloid LT-HSC signature (*top*), and non-myeloid LT-HSC signature (*bottom*) from Mann et al., 2018 (24). NES, normalized enrichment score. (**E to G**) Barplots showing log_2_ and DESeq2-normalized gene expression for select genes associated with (**E**) granulocyte or monocyte, (**F**) platelet, or (**G**) stem programs. Statistical significance was calculated using a paired, two-tailed Student’s *t*-test adjusted for multiple hypothesis testing with Benjamini-Hochberg procedure. \**P-adjusted* < 0.05, \*\**P-adjusted* < 0.01, \*\*\**P-adjusted* < 0.001, \*\*\*\**P-adjusted* < 0.0001. All barplots in this figure indicate mean ± SEM.

We also searched for the expression of lineage-specific transcripts that may indicate signs of early myeloid and lymphoid priming in LT-HSCs. Among the genes significantly enriched in NEO1^+^ compared to NEO1^−^*Hoxb5*^+^ LT-HSCs, we found several myeloid genes, including *Lrg1* and lineage-related transcription factors, such as *Meis2*, *Hoxb6,* and *Cebpe* (Fig. 5*E*), and platelet genes (Fig. 5*F*), such as *Vwf*, *Clu*, and *Selp*. We also find that NEO1^+^ LT-HSCs are significantly enriched (*Q* < 0.05) for previously reported gene signatures of megakaryocyte progenitors (MkP) and pre-erythrocyte colony-forming units (preCFU-E) (*SI Appendix,* Fig. S7*C*) (27). Moreover, the gene expression signature of NEO1^+^*Hoxb5*^+^ LT-HSCs significantly aligned with expression profiles of previously reported myeloid-biased LT-HSCs (Fig. 5*D***, *top***), while NEO1^−^ LT-HSCs were enriched for the balanced LT-HSC signature (Fig. 5*D*, ***bottom***). Altogether, these data suggest that LT-HSCs may sample regions of the transcriptome associated with their lineage fate decisions.

Finally, we also compared NEO1^+^ and NEO1^−^*Hoxb5*^+^ LT-HSC gene expression with respect to known stemness-associated genes. Overall, NEO1^+^*Hoxb5*^+^ LT-HSCs had higher expression of known stem-related genes, including *Ctnnal1* (14), *Fgd5* (13), *Bmi1* (54), *Gprc5c* (39), and *Slamf1* (11) (Fig. 5*G*), the latter of which was confirmed by flow cytometry (*SI Appendix,* Fig. S8). This suggests that although previously identified stemness genes enrich for a self-renewing phenotype, their expression may also be associated with myeloid-bias.

### Lineage-balanced NEO1^−^*Hoxb5*^+^ LT-HSCs outcompete NEO1^+^*Hoxb5*^+^ LT-HSCs in reconstitution and reside at the apex of hematopoiesis

To directly compare the relative fitness of NEO1^+^*Hoxb5*^+^ LT-HSCs and NEO1^−^*Hoxb5*^+^ LT-HSCs, we co-transplanted 200 cells from each fraction with host supporter cells into irradiated 2-to-3-month-old congenic recipients (Fig. 6*A*). Donor origin was distinguished by CD45.2 and EGFP expression using Hoxb5-mCherry and *CAG*-EGFP;*Hoxb5*-mCherry mice (Fig. 6*A*). Unlike the primary transplants above, NEO1^−^*Hoxb5*^+^ LT-HSCs exhibited significantly higher reconstitution potential compared to NEO1^+^*Hoxb5*^+^ LT-HSCs in the co-transplantation setting (Fig. 6*B*). This suggests that NEO1^−^*Hoxb5*^+^ LT-HSCs are more fit to reconstitute young recipients compared to NEO1^+^*Hoxb5*^+^ LT-HSCs.

**Figure 6.**
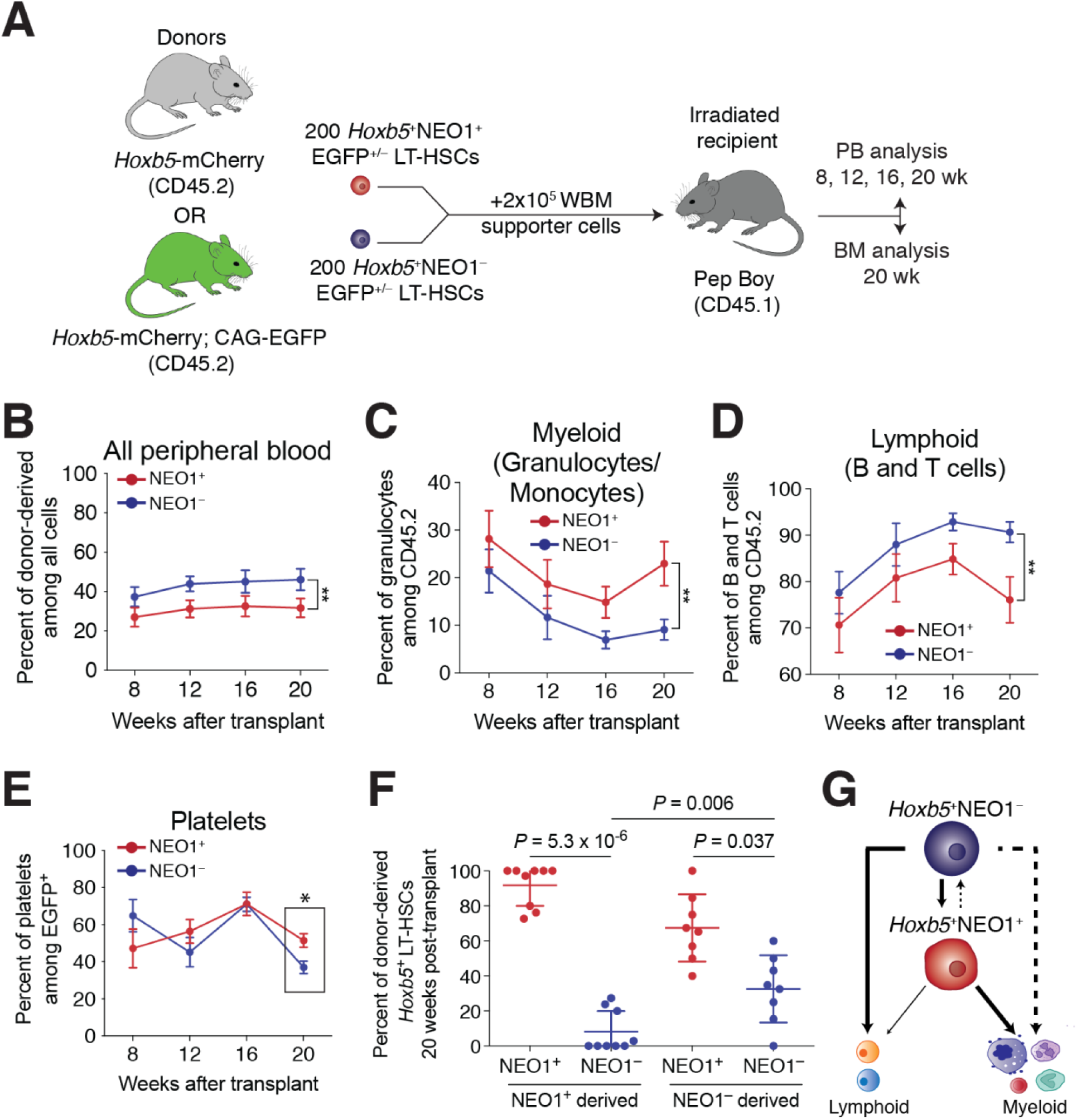
NEO1^−^*Hoxb5*^+^ LT-HSCs outcompete NEO1^+^*Hoxb5*^+^ LT-HSCs in reconstitution potential and reside at the apex of the hematopoietic hierarchy. (**A**) Experimental design for co-transplantation of NEO1^+^ and NEO1^−^*Hoxb5*^+^ LT-HSCs (*n* = 12 mice). (**B to D**) Measurements of reconstitution potential and lineage priming after co-transplantation. (**B**) Percent of donor-derived cells among all peripheral blood cells at 8, 12, 16, and 20 weeks post-transplant. (**C**) Percent of myeloid cells (GR1^+^CD11B^+^ granulocytes and monocytes) among donor-derived cells at 8, 12, 16, and 20 weeks post-transplant. (**D**) Percent of lymphoid cells (B220^+^ B cells and CD3^+^ T cells) among donor-derived cells at 8, 12, 16, and 20 weeks post-transplant. Statistical significance for **B to D** was calculated using two-way ANOVA with time post-transplant and NEO1 status as factors. \*\**P* < 0.01. (**E**) Percent of platelets (EGFP^+^CD41^+^) among donor-derived cells at 8, 12, 16, and 20 weeks post-transplant. Statistical significance at 20 weeks post-transplant was calculated using an unpaired, two-tailed Student’s *t*-test. \**P* < 0.05. (**F**) Percent of NEO1^+^ and NEO1^−^ *Hoxb5*^+^ LT-HSCs derived from donor NEO1^+^ and NEO1^−^*Hoxb5*^+^ LT-HSCs in the mouse bone marrow 20 weeks post-transplant. Only samples for which *Hoxb5*^+^ LT-HSCs were present are shown (‘NEO1^+^ derived’, *n* = 9; ‘NEO1^−^ derived’, *n* = 8). Statistical significance was calculated using a paired, two-tailed Student’s *t*-test between the percent of NEO1^+^ and NEO1^−^ *Hoxb5*^+^ LT-HSCs derived from the same donor and an unpaired, two-tailed Student’s *t*-test between the percent of NEO1^−^*Hoxb5*^+^ LT-HSCs between NEO1^+^ and NEO1^−^ donors. *P* values are indicated on the graph. (**G**). Schema depicting a revised model of long-term hematopoiesis with a lineage-balanced, quiescent NEO1^−^*Hoxb5*^+^ LT-HSC residing above a downstream myeloid-biased NEO1^+^*Hoxb5*^+^ LT-HSC. The thickness of the lines indicates the degree of contribution and dashed lines mark putative differentiation paths. All line plots in this figure indicate mean ± SEM. Scatter dot plot in **F** indicates mean ± SD.

Co-transplantation also confirmed that NEO1^+^*Hoxb5*^+^ LT-HSCs contribute significantly more to granulocytes and monocytes (myeloid) and less to B and T cells (lymphoid) compared to NEO1^−^ *Hoxb5*^+^ LT-HSCs (Fig. 6 *C* and *D*). We also quantified platelet fractions among EGFP^+^ donors. Relative platelet contribution from NEO1^+^*Hoxb5*^+^ was not significantly different from NEO1^−^ *Hoxb5*^+^ donors during the first 16 weeks post-transplant, but significantly increased among NEO1^+^*Hoxb5*^+^-derived PB at 20 weeks post-transplant (Fig. 6*E*).

Finally, we also measured the composition of LT-HSCs between NEO1^+^ and NEO1^−^ derived bone marrow in the co-transplantation setting. Both NEO1^+^ and NEO1^−^*Hoxb5*^+^ cells produced on average equal numbers of *Hoxb5*^+^ LT-HSCs per million bone marrow cells (*SI Appendix,* Fig. S9), and among *Hoxb5*^+^ LT-HSCs, more NEO1^+^ than NEO1^−^*Hoxb5*^+^ cells (Fig. 6*F*). However, NEO1^+^*Hoxb5*^+^ cells produced significantly fewer NEO1^−^*Hoxb5*^+^ cells compared to NEO1^−^ *Hoxb5*^+^ cells (*P* = 0.006). This suggests limited transition from NEO1^+^ to NEO1^−^, while NEO1^−^ *Hoxb5*^+^ cells are capable of giving rise to high percentages of both populations (Fig. 6*F*). Therefore, NEO1^−^*Hoxb5*^+^ LT-HSCs likely precede NEO1^+^*Hoxb5*^+^ LT-HSCs in the differentiation hierarchy (Fig. 6*G*).

## DISCUSSION

Previously, we showed that phenotypic HSCs (pHSCs) are variable in their reconstitution potential and that *Hoxb5* expression distinguishes long-term from short-term repopulating HSCs (6). While *Hoxb5-* pHSCs were unable to repopulate secondary recipients and were homogeneously lymphoid-biased, Hoxb5+ pHSCs serially reconstituted recipients and exhibited variable contribution to hematopoietic lineages. Therefore, we sought to understand the diversity of self-renewing HSCs in the mouse bone marrow using *Hoxb5* as a reporter to mark long-term HSCs (LT-HSCs).

To accomplish this, we screened gene expression profiles for candidate surface markers that are strictly enriched in HSCs and stratify *Hoxb5*^+^ LT-HSCs into subpopulations. We identified Neogenin-1 (*Neo1*; NEO1) as a transmembrane receptor specifically expressed on a sub-fraction of *Hoxb5*^+^ LT-HSCs. Although NEO1 is well known for its role in the brain (29–31), skeletal muscle (32), breast epithelia (34), endothelia (35), and other tissues (33), its expression in the bone marrow and association with LT-HSCs has not yet been extensively studied. We find that NEO1^+^ cells comprise a minor fraction of *Hoxb5*^+^ LT-HSCs in young mice that progressively expands with age and represents >80% of 22-month-old *Hoxb5*^+^ LT-HSCs. This expansion can be partially explained by the higher frequency of NEO1^+^*Hoxb5*^+^ in the G_2_/S cell cycle phase compared to NEO1^−^*Hoxb5*^+^ LT-HSCs. Both NEO1^+^ and NEO1^−^*Hoxb5*^+^ LT-HSCs are long-term self-renewing and contribute to all major blood lineages during primary and secondary transplantations into irradiated mice. However, NEO1^+^*Hoxb5*^+^ LT-HSCs exhibit a stable bias towards myeloid lineages and are less productive in secondary transplants compared to NEO1^−^*Hoxb5*^+^ LT-HSCs. Gene expression comparison reveals higher cell cycle and lower ribosomal gene expression in NEO1^+^ than NEO1^−^ LT-HSCs. This is consistent with the higher incidence of NEO1^+^*Hoxb5*^+^ LT-HSCs in the G_2_/S than G_1_ cell cycle phase (55). NEO1^+^*Hoxb5*^+^ LT-HSCs also exhibited early sampling of myeloid- and platelet-related genes, including lineage-related transcription factors, at the transcript-level.

Several previous studies have used single cell transplants to describe HSC heterogeneity and the existence of different lineage-primed states. Single cell HSC transplants by Dykstra and colleagues described two fractions of long-term self-renewing HSCs, ‘α cells’ and ‘β cells’, which were myeloid-biased and lineage-balanced, respectively (1). Yamamoto and colleagues also have demonstrated the presence of myeloid-restricted progenitors with long-term repopulating activity (MyRPs) that are derived directly from balanced HSCs and expand with age (2, 3). While single HSC transplants evidenced the presence of balanced and myeloid-biased LT-HSCs, these studies did not identify surface or transcriptional markers to distinguish these populations. We have demonstrated a strategy for the prospective isolation of balanced and myeloid-biased LT-HSCs using Neogenin-1 and *Hoxb5*. The functional potential of NEO1^+^ and NEO1^−^*Hoxb5*^+^ LT-HSCs matches the characteristics of the myeloid-biased and balanced LT-HSC fractions previously predicted by single cell transplants (1–3).

Other markers have also been proposed to enrich for myeloid-biased HSCs. For example, our group previously showed that high CD150 surface expression enriches for myeloid-biased HSCs (21). However, the association of CD150 with myeloid-bias was not evaluated in long-term repopulating cells and additional marker combinations can improve the purification of balanced from myeloid-biased HSCs. Nevertheless, *Hoxb5*^+^NEO1^+^ LT-HSCs indeed express higher CD150 (*SI Appendix,* Fig. S7), validating our initial attempts to prospectively isolate myeloid-biased HSCs (21). CD41 has also been suggested to mark myelo-erythroid HSCs (26), although it likely separates different fractions. CD41^−^ HSCs are lymphoid biased and proliferative, while NEO1^−^*Hoxb5*^+^HSCs are balanced and quiescent. The vWF reporter mouse is another system used to isolate platelet-biased and myeloid-biased HSCs (27). However, like the CD41^−^ HSCs, vWF^−^ HSCs are lymphoid-biased HSCs that are phenotypically distinct from the balanced, quiescent NEO1^−^*Hoxb5*^+^ cells we describe in this study. Finally, CD61 was recently described as a surface marker on long-term repopulating myeloid-biased LT-HSCs that respond to inflammatory stress and expand with age (24). The CD61^+^ and CD61^−^ LT-HSCs are transcriptionally similar to NEO1^+^ and NEO1^−^*Hoxb5*^+^ LT-HSCs, suggesting that these markers may capture similar cell types (Fig 5 *Ea* and *F*). Leveraging combinations of both surface markers will likely improve the purification of balanced LT-HSCs from lineage-biased LT-HSCs.

Our results also bring to question the hierarchical relationship between lineage-primed and balanced LT-HSCs. Previous single cell transplants studies suggest some degree of plasticity between LT-HSC fractions with different differentiation potentials (1). Our co-transplantation experiment indicates that NEO1^−^*Hoxb5*^+^ LT-HSCs are likely upstream of NEO1^+^*Hoxb5*^+^ LT-HSCs, as NEO1^−^*Hoxb5*^+^ LT-HSCs produced NEO1^+^ *Hoxb5*^+^ LT-HSCs, while conversion of NEO1^+^ *Hoxb5*^+^ LT-HSCs to NEO1^−^*Hoxb5*^+^ HSCs was rare. The few instances of NEO1^+^ donors producing NEO1^−^*Hoxb5*^+^ LT-HSCs may be attributable to impurity among the 200 cells that were injected. This is also consistent with our gene expression and cell cycle analysis demonstrating that NEO1^+^*Hoxb5*^+^ LT-HSCs are more often cycling compared to NEO1^−^*Hoxb5*^+^ cells (56). Moreover, NEO1^−^*Hoxb5*^+^ LT-HSCs contributed more to total hematopoiesis in secondary transplants and co-transplants compared to NEO1^+^*Hoxb5*^+^ LT-HSCs. Therefore, our data suggest that balanced quiescent LT-HSCs reside at the apex of the hematopoietic hierarchy, corroborating a recent study showing that myeloid-biased MyRPs are derived from balanced HSCs (2, 3). However, this contrasts with the view that vWF^+^ and CD41^+^ platelet/myeloid-biased cells reside at the apex of the hematopoietic hierarchy (26, 27).

Additional work will also be required to delineate the differentiation path that distinct LT-HSC fractions follow to generate various blood cells. Our study suggests that balanced NEO1^−^*Hoxb5*^+^ LT-HSCs contribute to myeloid lineage through a NEO1^+^*Hoxb5*^+^ LT-HSC intermediate. However, it is unclear whether NEO1^−^*Hoxb5*^+^ LT-HSCs require the NEO1^+^*Hoxb5*^+^ intermediate state or can independently produce lineage progenitors through alternative routes. Moreover, it remains to be answered whether all NEO1^+^*Hoxb5*^+^ LT-HSCs are derived from NEO1^−^*Hoxb5*^+^ LT-HSCs. To fully reveal the hierarchical order and sequence of differentiation events, single cell tracking experiments will be required. Critically, our experimental results are based on the behavior of cells upon transplantation into young, irradiated mice. A recent study using individually barcoded HSCs showed that lineage biases are more pronounced after transplantation into lethally irradiated mice compared to unirradiated or anti-c-KIT-depleted syngeneic mice (4). This suggests that post-transplant lineage bias may be either due to plasticity in lineage output or the selective engraftment of pre-existing HSC subsets. Therefore, it will be important to evaluate the potential and hierarchical relationship between NEO1^+^ and NEO1^−^*Hoxb5*^+^ LT-HSCs during *in situ,* unperturbed hematopoiesis with *in vivo* lineage tracing.

Furthermore, we note that comparing NEO1^+^ and NEO1^−^ fractions within pHSCs without gating *Hoxb5*^+^ LT-HSCs may mislead the significance of NEO1 to separate myeloid-biased from balanced cells. The vast majority of NEO1^−^ cells among pHSCs are short-term, lymphoid-biased *Hoxb5*^−^ ST-HSCs (6) that far outnumber the long-term, balanced NEO1^−^*Hoxb5*^+^ LT-HSCs we find in this study. NEO1 is also rarely expressed in *Hoxb5*^−^ downstream cells and bone marrow niche cells and its functional role in these populations has not been evaluated in this study.

The functional differences between NEO1^+^ and NEO1^−^ Hoxb5+ LT-HSCs may be influenced by intrinsic programs, external cues, or both. As NEO1 is a receptor to many known ligands, ongoing studies are evaluating the role of the bone marrow niche and particular ligands to NEO1 in influencing lineage bias and stem cell maintenance.

Finally, our antibody against NEO1 also identified higher proportions of cells in human HSCs compared to MPPs, LMPPs, and downstream progenitors, suggesting a possibly conserved role of NEO1 in human HSC biology. Therefore, further evaluation of NEO1^+^*Hoxb5*^+^ LT-HSCs and the receptor-ligand interactions may offer insights into evolutionarily conserved mechanisms of lineage bias during long-term hematopoiesis.

Taken together, we have identified a novel marker on the surface of *Hoxb5*^+^ LT-HSCs, Neogenin-1, that enables the separation of myeloid-biased LT-HSCs from quiescent, balanced LT-HSCs with the highest long-term repopulation potential. Our findings reveal a previously undescribed layer of functional heterogeneity among strictly defined functional LT-HSCs and enable the precise and prospective study of LT-HSCs and their fractions.

## MATERIALS AND METHODS

### Mice

2-to-3-month-old female *Hoxb5*-mCherry mice (MGI:5911679; available through RIKEN BioResource Research Center: RBRC09733) were used as donors (CD45.2) for transplant experiments and bulk RNA-sequencing. Additionally, 2-to-3-month-old *CAG-EGFP*;*Hoxb5*-mCherry mice (in-house colony) were used for co-transplant assays. 2-to-3-month-old female B6.SJL-*Ptprc*^a^ *Pepc*^b^/BoyJ mice (Jackson Laboratory) were used as recipients (CD45.1) for transplant experiments and for supporter bone marrow. 4-month-old female *Hoxb5*-mCherry mice were used for experiments with 5-fluorouracil. 5-month, 13-month, and 22-month-old female *Hoxb5*-mCherry mice were used for aging analysis. 2-to-3-month-old and 12-to-14-month-old female *Hoxb5*-mCherry mice were used for cell cycle analysis. 2-to-3-month-old C57BL/6J female mice (Jackson Laboratory) were used for fluorescent-minus-one (FMO) controls for *Hoxb5*-mCherry expression.

### Gene expression profiles of mouse hematopoietic cells and the bone marrow niche

All microarray data used in this study are accessible through the Gene Expression Commons platform (http://gexc.stanford.edu) and the Gene Expression Omnibus (GEO) accession, GSE34723. We analyzed 64 microarray gene expression profiles (GEPs) of 23 distinct mouse hematopoietic cell types (*SI Appendix,* Table S1) for surface markers enriched in HSCs compared to downstream progeny. GEPs were normalized against a large common reference of >11,939 Affymetrix Mouse Genome 430 2.0 microarrays as described before(36). For each gene, the probeset with the largest dynamic range was selected and transformed to percentile ranks (range: −100% to +100%) based on its relative expression to the reference.

Genes were further subset based on two main criteria (1) positive percentile expression in HSCs and (2) annotation as a cell surface protein based on GO:0009986, leaving 186 gene candidates. Fold-change enrichment in HSCs was calculated as the average percentile rank for each gene among HSCs divided by the average percentile rank for that gene across all other cells.

Single cell transcriptomes of hematopoietic stem and progenitors (HSPCs) were acquired from GEO accessions, GSE90742, and from the Single-Cell Gene Expression Atlas for hematopoietic cells (http://blood.stemcells.cam.ac.uk/single_cell_atlas.html). In the dataset from Rodriguez-Fraticelli et al. (5), cells with greater than 0 unique molecular identifiers (UMIs) were considered *Neo1*^+^. In the dataset from Nestorowa et al. (57), cells with greater than 4 log_2_ counts were considered *Neo1*^+^. A threshold of 4 was chosen based on the color gradient thresholds for the diffusion maps in the Single-Cell Gene Expression Atlas.

tSNE plots of single cell RNA-sequencing data from the mouse bone marrow niche were generated using the online tool nichExplorer (https://compbio.nyumc.org/niche/)(58).

### Bone marrow imaging

Femurs were collected and fixed in 4% PFA with 10% EDTA solution at 4°C overnight. After decalcification in 20% EDTA solution at 4°C for two days, bones were saturated in increasing concentrations of sucrose solution (10% for 1h, 20% for 1h, and 30% overnight) at 4°C. Prepared bones were frozen in OCT compound using an isopentane bath cooled on dry ice. For staining, 50-70 µm thick sections were cut on microscope slides and blocked in 3% BSA with 10% FBS solution (blocking buffer) for 1h at room temperature. Sections were stained with primary antibodies against NEO1 (R&D cat. no. AF1079; dilution: 1:100) and Endomucin (ThermoFisher Scientific; clone: V.7C7; dilution: 1:100) in 10% blocking buffer at 4°C overnight and washed with PBS. For secondary staining, sections were incubated with donkey anti-rat IgG (H+L) AF488-conjugated and donkey anti-goat IgG (H+L) Cy3-conjugated (Jackson ImmunoResearch; dilution: 1:400) in 10% blocking buffer at room temperature for 1h. After another wash with PBS, samples were counterstained with DAPI and sealed for imaging. Images were acquired using a LSM880 confocal microscope (Zeiss) with LD LCI Plan-Apochromat 25x/0.8 objective. Images were analyzed using the Fiji software.

### Transplantation assays

2-to-3-month-old female B6.SJL-*Ptprc^a^ Pepc^b^* /BoyJ (CD45.1) recipient mice were lethally irradiated at a single dose of 9 Gy. For reconstitution assays, 10 NEO1^+^*Hoxb5*^+^ or NEO1^−^ *Hoxb5*^+^ LT-HSCs were isolated from donor CD45.2^+^ *Hoxb5*-mCherry mice (MGI:5911679) as described in sections *‘Bone marrow isolation’* and *‘Flow cytometry’* and co-injected with 2×10^5^ recipient whole bone marrow cells in 200 μl of PBS with 2% FBS into the retro-orbital venous plexus. For secondary transplants, 1000 CD45.2^+^ Lin^−^cKIT^+^SCA1^+^ (KLS) cells were isolated by flow cytometry and transplanted together with 2 x 10^5^ recipient (CD45.1) whole bone marrow cells into lethally irradiated recipient CD45.1^+^ mice as described above. For co-transplantation assays, 200 NEO1^+^ and NEO1^−^*Hoxb5*^+^ LT-HSCs were isolated from either CD45.2^+^ *Hoxb5*-mCherry mice (MGI:5911679) or an in-house strain of EGFP^+^CD45.2^+^ *Hoxb5*-mCherry mice and transplanted into lethally irradiated recipient CD45.1^+^ mice at a split dose of 9 Gy with a 4-hour interval, controlling for donor strain biases by transplanting the same condition from both strains. In all cases, recipients with lower than 1% total chimerism were considered failed transplantations and excluded from analysis.

### Peripheral blood analysis for chimerism

Peripheral blood collections for assessment of donor chimerism were performed at 4, 8, 12, and 16 weeks after primary and secondary transplantations and 8, 12, 16, and 20 weeks after co-transplantations. At each time point, 50-100 μl of blood was collected from the retro-orbital venous plexus using heparinized capillary tubes (Fisher Scientific) and added to K_2_/EDTA-coated MiniCollect tubes (Greiner Bio-One). Red blood cells were depleted with two rounds of ACK lysis buffer by incubating at RT for 5 min each. Cells were then washed with cold PBS. Cells were Fc-blocked with of rat IgG (LifeSpan BioSciences) and stained with 5 μg/ml of rat anti-mouse antibodies (catalog no., concentrations, and clone provided in Table S3 of the *SI Appendix*) to CD45.1, CD45.2, CD11B, GR1, B220, CD3, and only in the co-transplantation assay, CD41. 7-Aminoactinomycin D (7-AAD; BD Bioscience) was added for live and dead cell discrimination.

For reconstitution assays, total donor chimerism was defined as the percentage of CD45.1^−^ CD45.2^+^ cells among total CD45.1^−^CD45.2^+^ and CD45.1^+^CD45.2^−^ cells. For co-transplantation assays, total donor chimerism was defined as the percentage of either CD45.1^−^CD45.2^+^EGFP^−^ cells or CD45.1^−^CD45.2^+^EGFP^+^ cells among total CD45.1^−^CD45.2^+^EGFP^−^, CD45.1^−^ CD45.2^+^EGFP^+^, and CD45.1^+^CD45.2^−^EGFP^−^ cells. For all cases, lineage chimerism was evaluated as the percentage of lymphoid (B220^+^ B cells and CD3^+^ T cells) or myeloid (GR1^+^CD11B^+^ granulocytes and monocytes among donor-derived cells. Only EGFP^+^ cells were evaluated for platelet chimerism. In these cases, platelet chimerism was calculated as the percent of CD41^+^ platelets among all donor-derived EGFP^+^ cells and platelets.

### Mouse hematopoietic stem cell isolation by flow cytometry

HSCs were isolated from 2-to-3-month-old, 5-month-old, 13-month-old, and 22-month-old bone marrow of *Hoxb5*-mCherry mice (MGI:5911679). For bone marrow isolation, tibia, femur, and pelvis were dissected, crushed with mortar and pestle in FACS buffer (2% fetal bovine serum (FBS) in PBS with 100 U/ml DNase), and the supernatant was collected. For WBM isolation, red blood cells were depleted with ACK lysis buffer by incubating at RT for 10 min and Fc-blocked by incubating with rat IgG (LifeSpan BioSciences) for 10 min. For c-KIT^+^ cell isolation, samples were Fc-blocked with rat IgG for 10 min, incubated in c-KIT magnetic beads (Miltenyi) with 100 U/ml DNase, and MACS-isolated using LS magnetic columns (Miltenyi) as per manufacturer’s protocol. For the cell cycle analysis, samples were blocked with TruStain FcX (CD16/32) instead of rat IgG, and enriched for c-KIT^+^ cells. Each sample was normalized to an equal number of cells (4.5 million c-KIT^+^ cells) and processed following a previously published protocol for cell cycle analysis of HSCs (59). Sample collections with fewer than 10 total *Hoxb5*^+^ LT-HSCs after flow cytometry analysis were considered underpowered for determining the distribution of cells in G_0_, G_1_, and G_2_/S and therefore excluded.

Samples for mouse HSC isolation were stained with a cocktail of antibodies against lineage markers, i.e. CD3, Gr-1, CD11B, B220, and TER119 (AF700), c-KIT (APC-Cy7), SCA-1 (PE-Cy7), CD48 (BV711), FLK2 (PerCP-Cy5.5), CD150 (BV421), CD34 (primary: biotin; secondary: Strep-BUV737), and NEO1 (primary: goat anti-mouse/human (R&D cat. no. AF1079); secondary: donkey anti-goat IgG (H+L) cross-absorbed AF488-conjugated; negative control: normal goat IgG). Although the antibody to NEO1 is polyclonal, results were consistent across multiple reagent lots and experiments. Antibody-labeled cells were shown to express higher NEO1 mRNA compared to unlabeled cells (*SI Appendix,* Fig. S7*A*).

Primary and secondary antibody incubations were 20-30 min each with 5 min wash step in between. Catalog number, concentrations, and clone information are provided in Table S3 of *SI Appendix*.

Flow cytometry and cell sorting were performed on the BD FACSAria and BD LSRFortessa. Gating strategy for the different populations is shown in *SI Appendix,* Fig. S3. 7-AAD or DAPI were used as a viability dye for dead cell exclusion, depending on the assay. All cells were suspended in FACS buffer (2% FBS in PBS) on ice unless otherwise indicated.

### Myeloablative stress with 5-fluorouracil (5-FU)

4-month-old female *Hoxb5*-mCherry mice were injected with 150 mg of 5-FU (Sigma-Aldrich) per kg body weight from a stock solution of 10 mg/ml in PBS (41). Bone marrow populations were isolated and analyzed 5 days after treatment as described above. Notably, given the upregulation of CD11B in HSCs post-treatment with 5-FU (41), the antibody to CD11B was omitted from the lineage staining panel.

### Human hematopoietic stem cell analysis by flow cytometry

Human CD34^+^ bone marrow cells were purchased from AllCells, Inc and their use were approved by the Stanford University Institutional Review Boards. To analyze human hematopoietic stem cell and progenitor populations, CD34^+^ bone marrow cells were stained against lineage markers, i.e. CD3, CD4, CD8, CD11B, CD14, CD19, CD20, CD56, and GPA (PE-Cy5), stem and progenitor markers, i.e. CD34 (APC-Cy7), CD38 (APC), CD45RA (BV785), and CD90 (FITC), and NEO-1 (primary: goat anti-mouse/human (R&D cat. No. AF1079); secondary: donkey anti-goat IgG (H+L) Cy3-conjugated; negative control: normal goat IgG). Propidium iodide (PI; Sigma-Aldrich) was added for live and dead cell discrimination. Catalog number, concentrations, and clone information are provided in Table S3 of *SI Appendix*. Gating strategy for the different populations is shown in Figure 1*E*. Flow cytometry was performed as described above.

### RNA extraction and library preparation for bulk RNA sequencing

For RNA-sequencing experiments, 250-500 cells from two pooled mice per sample were sorted directly into 100 μL of lysis buffer (Buffer RL) and RNA was isolated with the Single Cell RNA Purification Kit (Norgen Biotek Corp.) according to the manufacturer’s protocol. RNA quality was measured by capillary electrophoresis using the Agilent 2100 Bioanalyzer with Nano mRNA assay at the Stanford Protein and Nucleic Acid (PAN) Facility.

Libraries were prepared using the Smart-seq2 protocol by Picelli et al., 2014 with minor modifications. Briefly, cDNA was generated by oligo-dT primed reverse transcription with MMLV reverse transcriptase (SMARTScribe, Clontech) and a locked template-switching oligonucleotide (TSO). This was followed by 18 cycles of PCR amplification using KAPA HiFi hotStart ReadyMix and ISPCR primers. Amplified cDNA was then purified using 0.7x volume Agencourt AMPure XP beads to remove smaller fragments. The resulting cDNA concentration and size distribution for each well was determined on a capillary electrophoresis-based Agilent 2100 Bioanalyzer with High Sensitivity DNA chip at the Stanford PAN facility. 40 ng of cDNA was then tagmented, uniquely barcoded, and PCR enriched using the Nextera DNA Library Prep Kit (Illumina, San Diego, CA). Libraries were then pooled in equimolar amounts and purified of smaller fragments using 0.7x Agencourt AMPure XP beads. Pooled libraries were checked for quality using the Agilent Bioanalyzer with High Sensitivity DNA chip at the Stanford PAN facility. 10 samples were sequenced with 151 bp paired-end reads on a single lane of NextSeq 500 (Illumina, San Diego, CA) at the Stanford Functional Genomics Facility.

After sequencing, bcl2fastq2 v2.18 (Illumina) was used to extract the data and generate FASTQ files for each sample by using unique barcode combinations from the Nextera preparation. Raw reads were trimmed for base call quality (PHRED score >=21) and for adapter sequences using Skewer v0.2.2 (60). Trimmed reads were then aligned to the mouse genome assembly (mm10) from UCSC (http://genome.ucsc.edu) using STAR v2.4 with default setting (61).

### RNA sequencing analysis

Count normalization and differential gene expression analysis was performed using the DESeq2 v1.22.2 package in R (45). Raw counts from STAR were inputted into a DESeqDataSet object indicating NEO1 status (‘status’) and mouse subject (‘subject’) as factors (‘design = ∼subject + status’). Counts were size-factor normalized using the ‘DESeq’ function and log_2_-transformed. Pairwise differential gene expression analysis was performed using the lfcShrink function and indicating ‘type = apeglm’, which applies the adaptive t prior shrinkage estimator. As recommended (45), a threshold of *P*-adjusted < 0.1 was used to define significance for differentially expressed genes (*SI Appendix,* Table S2).

Gene set enrichment analysis (GSEA) was performed using the GSEA software provided by the Broad Institute (51) and the clusterProfiler v3.10.0 (52) and HTSanalyzer v2.34.0 (62) packages in R. Hypergeometric test with GO: Biological Processes and dot plots in Figure S7 were generated using the clusterProfiler package in R. Gene expression signatures of myeloid and non-myeloid LT-HSCs were acquired from the original study by Mann et al., 2018 (24). Gene expression signatures of lineage-restricted progenitors, including megakaryocyte progenitors (MkP), pre-erythrocyte colony-forming units (preCFU-E), pre-granulocyte/macrophage progenitors (preGM), pre-megakaryocyte/erythrocyte progenitors (preMegE), and common lymphoid progenitors (CLP), were acquired from the original study by Sanjuan-Pla et al., 2013 (27).

### Statistics

Statistical significance between two groups was determined using a paired or unpaired Student’s *t* test, as appropriate. For comparison of two groups across multiple time points, statistical significance was determined using a two-way ANOVA using the groups and time points as factors. Multiple hypothesis correction was applied to gene expression comparisons using the Benjamini-Horchberg procedure. Results with *P-adjusted* < 0.1 were considered significant. Data analyses were performed with R 3.5.1, Prism v7 (GraphPad Software, Inc.), and FlowJo v10 (FlowJo, LLC). The investigators were not blinded to allocation during experiments and outcome assessment. No sample-size estimates were performed to ensure adequate power to detect a pre-specified effect size.

## Acknowledgements

We thank R. Yamamoto, C.K.F. Chan, J. Xiang, V. Mascetti, V.G. Alvarado, A. Chandra, and A. Manjunath for technical assistance and discussion. We are grateful to T. Naik for lab management, A. McCarty and C. Wang for mouse breeding and management, P. Lovelace and S. Weber for their support and assistance with FACS, and Stanford Functional Genomic Facility (SFGF) for assistance with sequencing scRNA-seq libraries. We thank K. Kao, T. Sakamaki, and M. Miyanishi for assistance with breeding the *Hoxb5-*mCherry and *CAG*-EGFP;*Hoxb5*-mCherry mouse strains.

## Author contributions

K.S. identified Neogenin-1. K.S., G.S.G., M.Z., and I.L.W. designed the experiments; K.S., G.S.G., M.Z., J.N., A.Z., R.S., B.G., and D.W. performed the experiments. K.S., G.S.G., M.Z., and J.N. analyzed the data. G.S.G., K.S., M.Z., and I.L.W. wrote the manuscript. I.L.W. supervised the study. All authors commented on the manuscript at all stages.

## Data and materials availability

All expression datasets from the public domain analyzed in this work are described in **Materials and Methods**. The bulk RNA-sequencing data generated in this study has been deposited in GEO under accession code GSE130504. All data generated or analyzed in this study have also been deposited in the Dryad Digital Repository under doi:10.5061/dryad.376gn3k.

## Supporting Information (SI) Appendix

Table S1 Gene expression percentiles of 64 microarray expression profiles from 23 distinct mouse hematopoietic cell types with surface marker annotations and fold enrichment values in HSCs

Table S2 Genes differentially expressed between NEO1^+^ and NEO1^−^ *Hoxb5*^+^ LT-HSCs by DESeq2

Table S3 Inventory of antibodies and reagents

**Figure S1.**
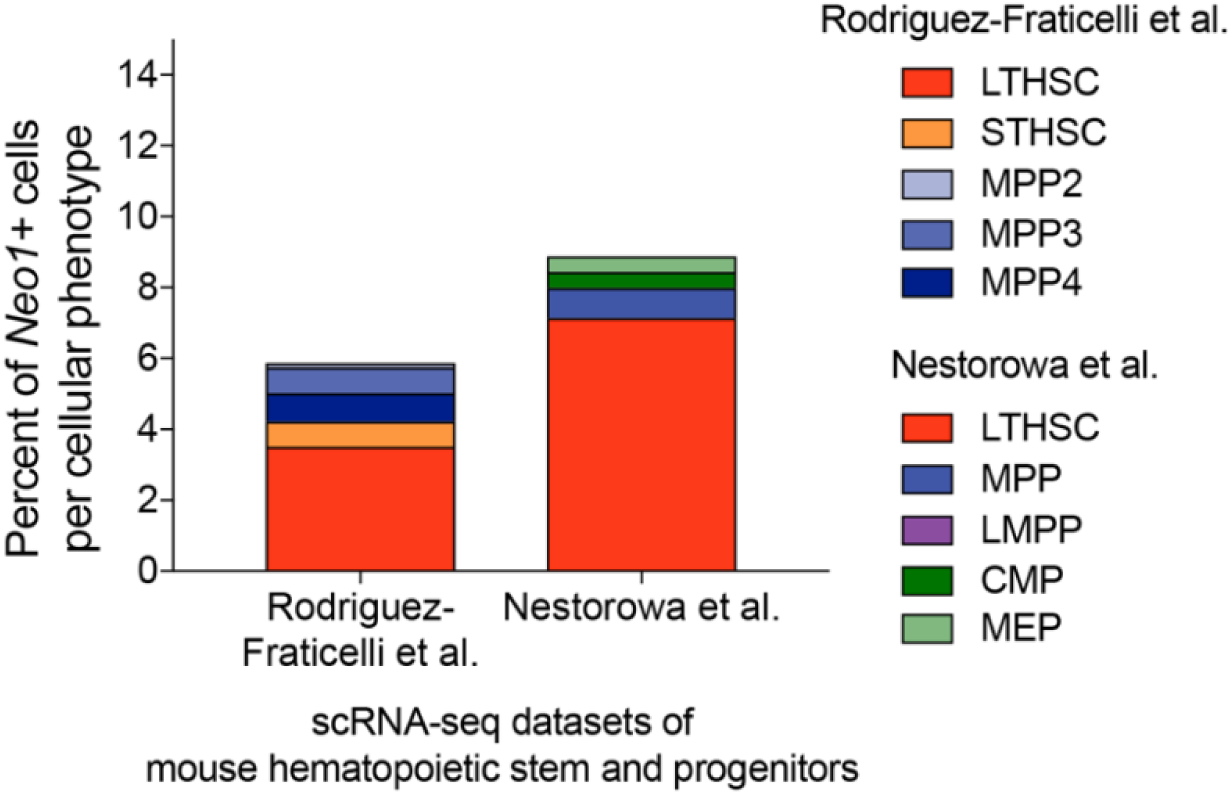
Single cell RNA sequencing shows selective expression of Neogenin-1 in a subset of LT-HSCs. Percent of *Neo1*^+^ cells among each phenotype defined by the authors of two independent scRNA-seq studies of mouse hematopoietic stem and progenitor cells (HSPCs) (5, 57).

**Figure S2.**
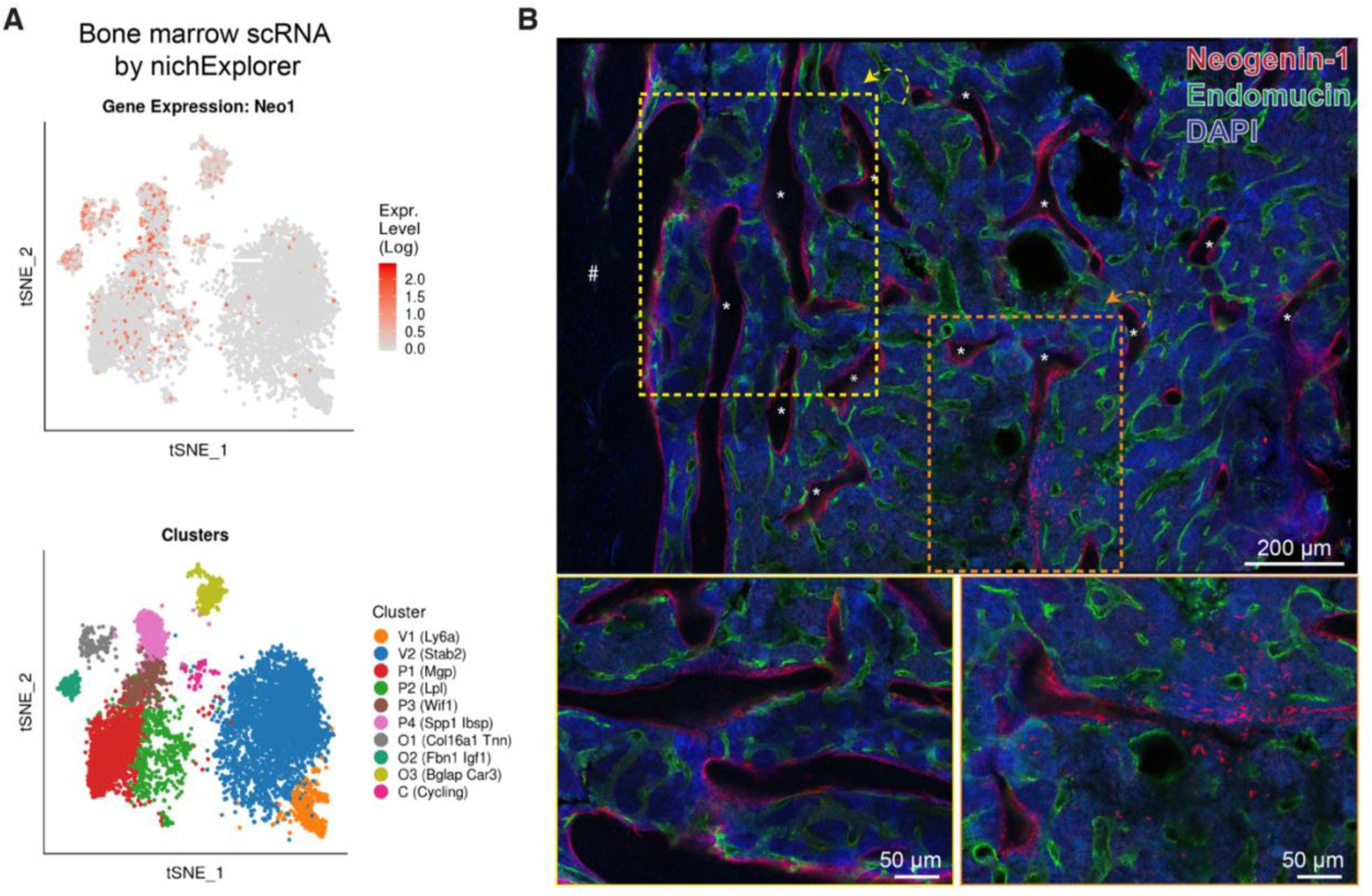
NEO1 expression in the bone marrow niche. (**A**) tSNE plot of *Neo1* expression (*top*) among different bone marrow niche populations (*bottom*) from a published scRNA-seq dataset of mouse osteoblasts and bone marrow stromal cells (58). (**B**) Representative images of bone marrow sections showing NEO1 (red) signal on the surface of trabecular bone (*) and stromal cells. Sinusoids are highlighted with an antibody to Endomucin (green) and nuclei were counterstained with DAPI (blue). “#” marks the bone surface. Scale bar is shown on the graph and boxes indicate corresponding magnified images below.

**Figure S3.**
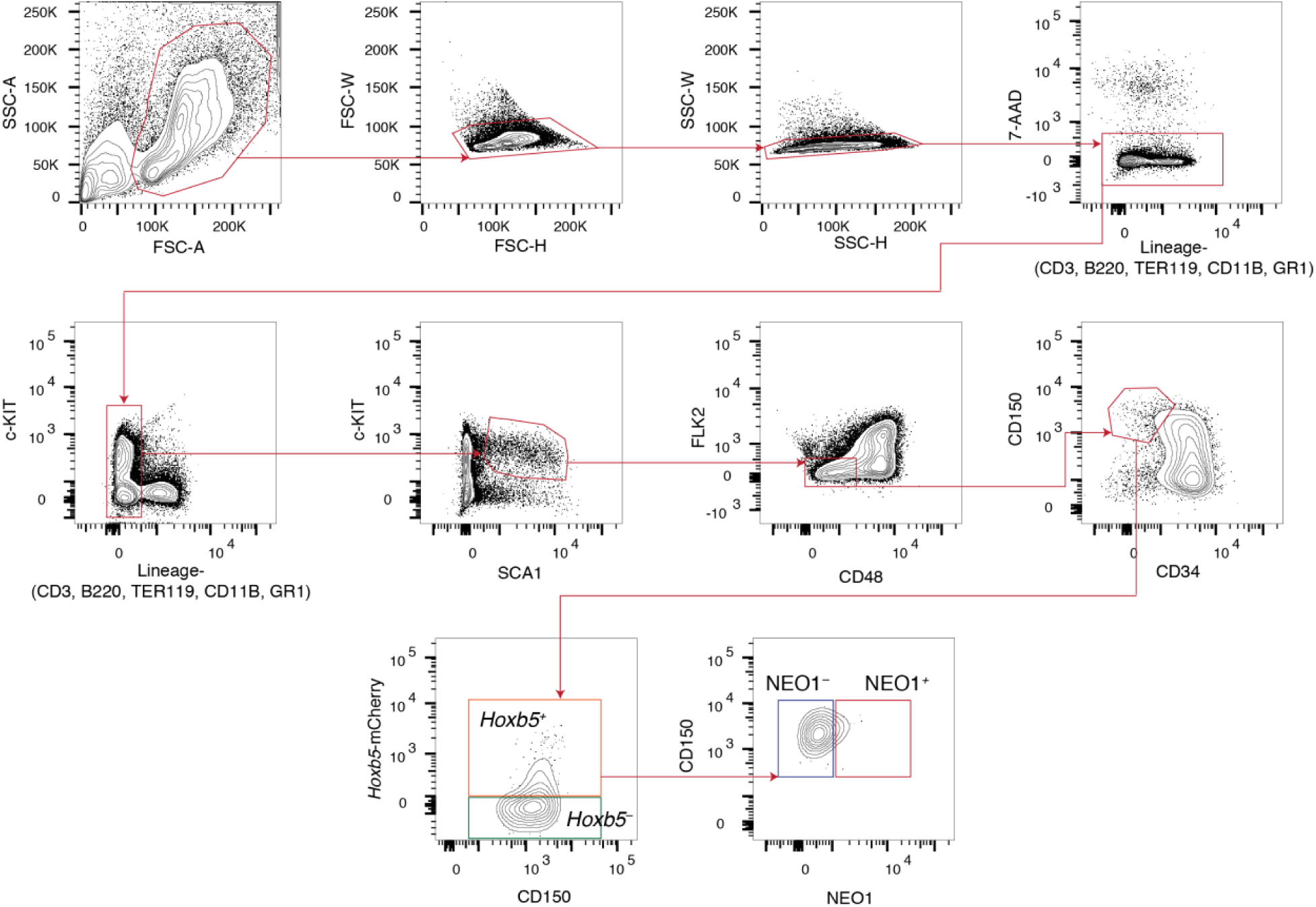
Gating scheme for the prospective isolation of NEO1^+^ and NEO1^−^ *Hoxb5*^+^ LT-HSCs by flow cytometry. Arrows indicate the gating sequence. All gates were drawn with respective to fluorescence-minus-one (FMO) controls. SSC, side scatter; FSC, forward scatter; 7-AAD, 7-Aminoactinomycin D.

**Figure S4.**
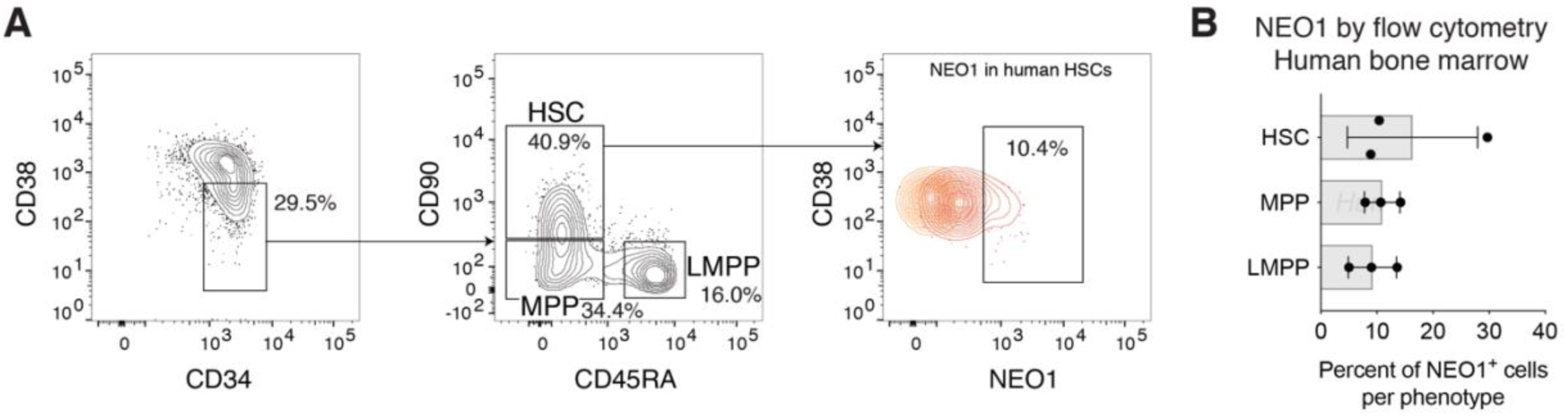
Flow cytometry analysis of NEO1 surface expression in the human bone marrow. (**A**) Contour plots with outliers showing the gating scheme for human HSCs, MPPs, and LMPPs (*n* = 3). (**B**) Barplots showing the percent of NEO1^+^ cells for each cell type gated in **A**. MPP, multipotent progenitor, LMPP, lymphoid-primed multipotent progenitor. Barplots indicate mean ± SD.

**Figure S5.**
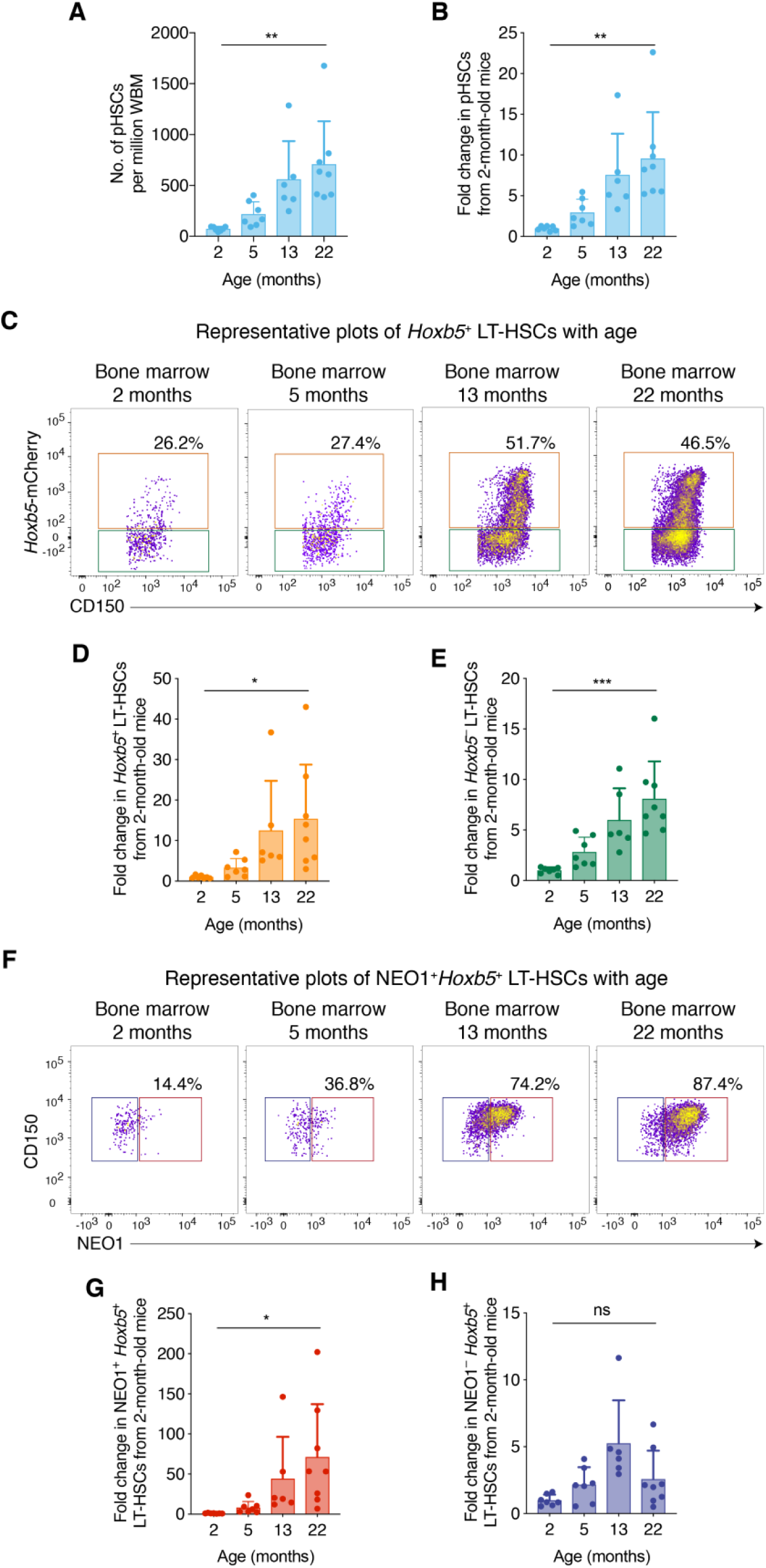
Fold-change in pHSCs, *Hoxb5*^−^ ST-HSCs, *Hoxb5*^+^ LT-HSCs, and NEO1^+^ and NEO1^−^ *Hoxb5*^+^ LT-HSCs with age. (**A and B**) Number and fold-change in pHSCs from mouse bone marrow at 2 (*n* = 7 mice), 5 (*n* = 7 mice), 13 (*n* = 6 mice), and 22 (*n* = 9 mice) months of age. Barplots showing (**A**) the number of pHSCs per million whole bone marrow (WBM) cells and (**B**) fold-change in pHSCs from 2-month-old mice. (**C**) Flow cytometry diagrams of *Hoxb5*-mCherry (*y*-axis) expression in the mouse bone marrow at 2 (*n* = 7 mice), 5 (*n* = 7 mice), 13 (*n* = 6 mice), and 22 (*n* = 9 mice) months of age. (**D and E**) Barplots showing the fold-change in (**D**) *Hoxb5*^+^ LT-HSCs and (**e**) *Hoxb5*^−^ ST-HSCs from 2-month-old mice. (**f**) Flow cytometry diagrams of NEO1 (*x*-axis) expression in the mouse bone marrow. (**G and H**) Barplots showing the fold-change in (**G**) NEO1^+^ *Hoxb5*^+^ LT-HSCs and (**H**) NEO1^−^ *Hoxb5*^+^ LT-HSCs from 2-month-old mice. For all barplots, statistical significance was calculated using an unpaired, two-tailed Student’s *t*-test between 2 months and 22 months of age. \**P* < 0.05, \*\**P* < 0.01, \*\*\**P* < 0.001, ns = non-significant, *P* ≥ 0.05. Bars and error bars indicate mean ± SD.

**Figure S6.**
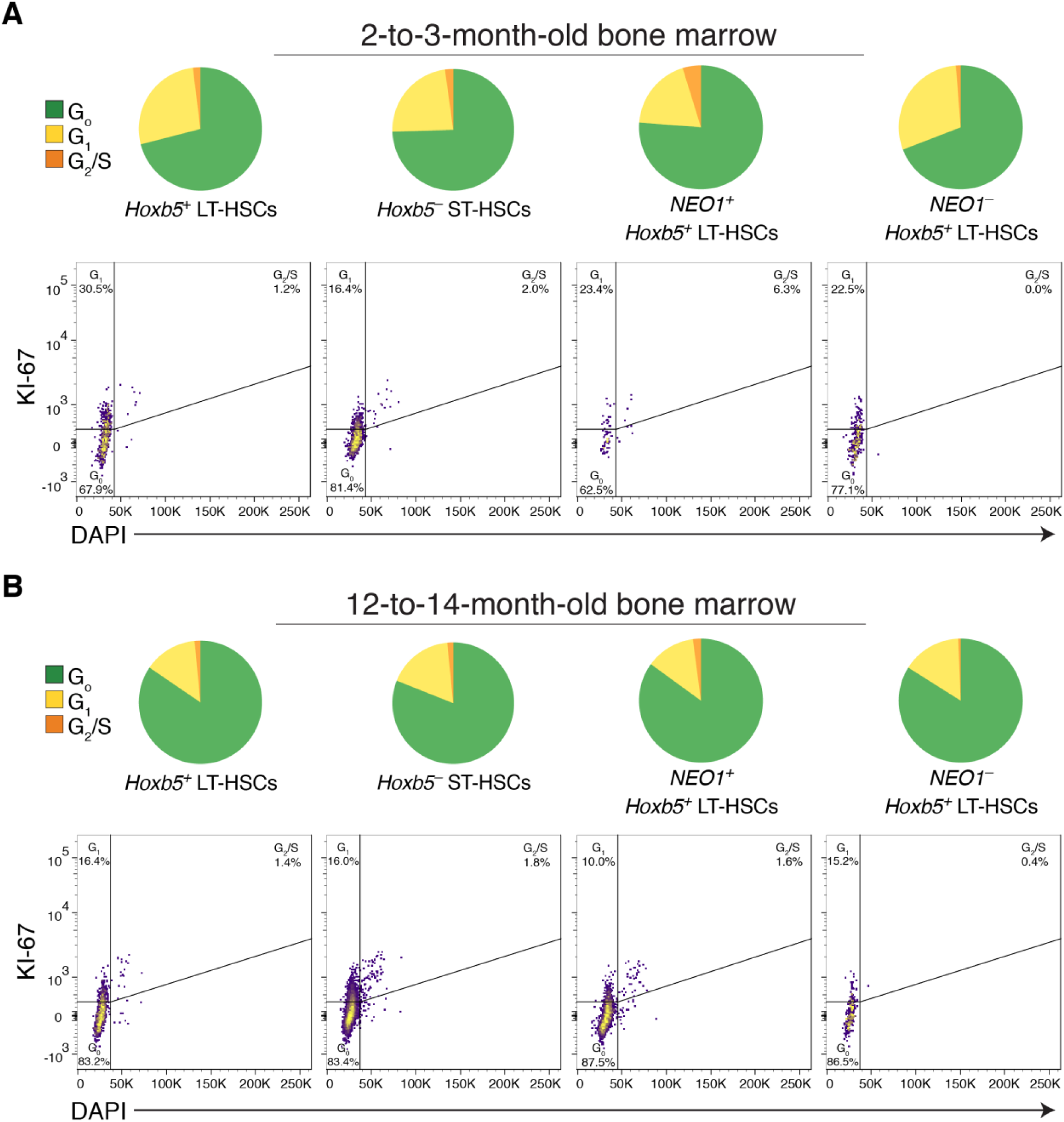
Cell cycle analysis of *Hoxb5*^−^ ST-HSCs*, Hoxb5*^+^ LT-HSCs, and NEO1^+^ and NEO1^−^ *Hoxb5*^+^ LT-HSCs. (**A**) Pie charts (*top*) showing the average percent of each cell type from 2-to-3-month-old mouse bone marrow in G_0_ (green), G_1_ (yellow), and G_2_/S (orange) as measured by flow cytometry analysis of KI-67 and DAPI staining (*bottom*). Same as in (**B**) but with 12-to-14-month-old mouse bone marrow. KI-67 gates were drawn with respective to a fluorescence-minus-one (FMO) control.

**Figure S7.**
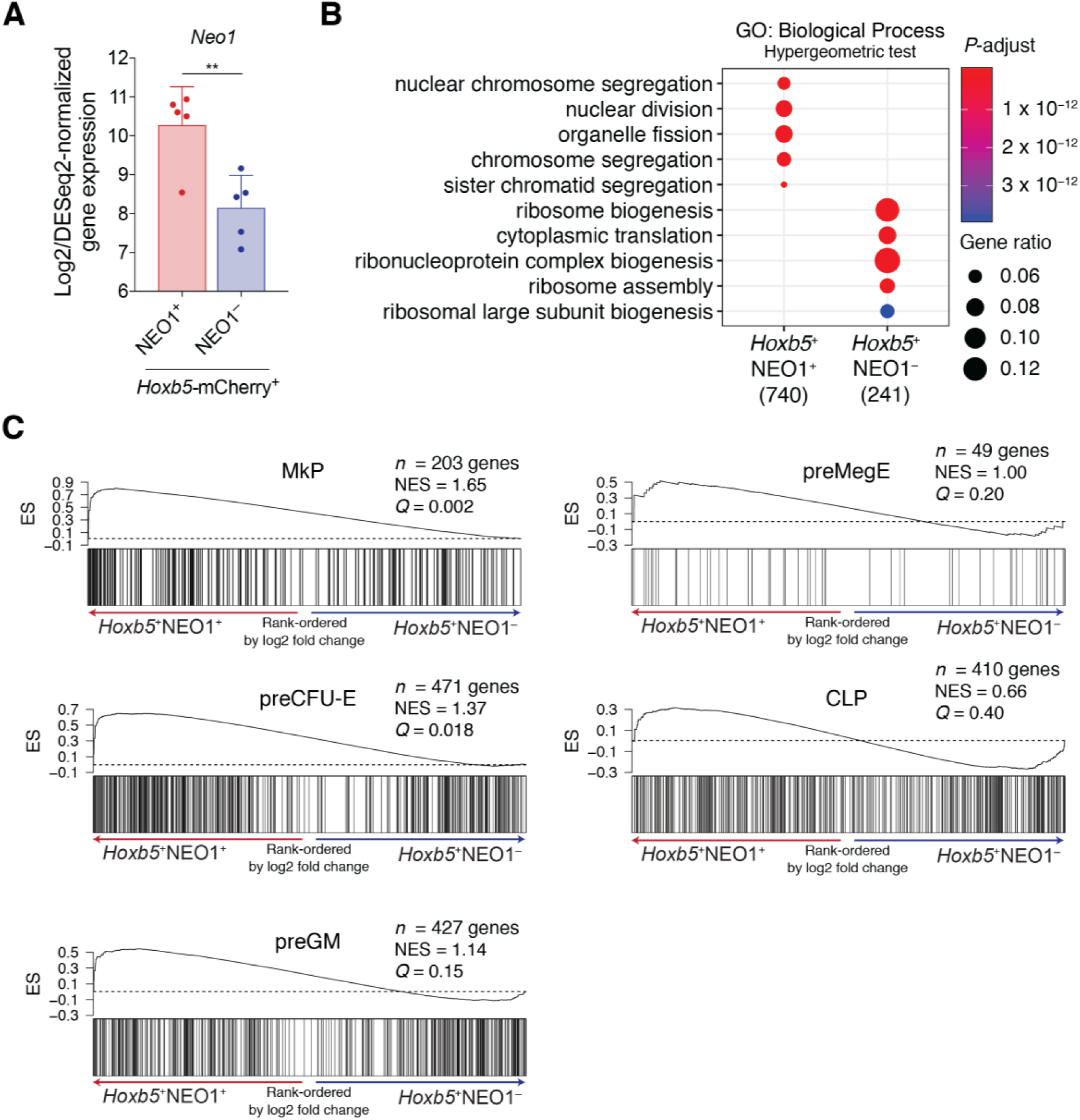
Enrichment of GO biological processes and lineage-restricted progenitor signatures in NEO1^+^ and NEO1^−^ *Hoxb5*^+^ LT-HSC transcriptomes. (**A**) Barplots showing log_2_ and DESeq2-normalized gene expression of *Neo1* in NEO1^+^ and NEO1^−^ cells isolated by flow cytometry. Statistical significance was calculated using a paired, two-tailed Student’s *t*-test adjusted for multiple hypothesis testing with Benjamini-Hochberg procedure. \*\**P* < 0.01. Bars and error bars indicate mean ± SEM. (**B**) Dot plots showing the top 5 ‘GO: Biological Process’ pathways enriched in NEO1^+^ and NEO1^−^ *Hoxb5*^+^ LT-HSCs by hypergeometric test. *P-*adjusted values are indicated by color gradient and gene ratios by dot size. Number of genes significantly enriched in NEO1^+^ or NEO1^−^ are indicated in parenthesis below each column. (**C**) GSEA plots showing enrichment of gene set signatures of lineage restricted progenitors from a previous study (27) over a gene list ordered by log_2_ fold change in NEO1^+^ versus NEO1^−^ *Hoxb5*^+^ LT-HSCs. NES, normalized enrichment score and *Q* values are indicated on the graph. MkP, megakaryocyte progenitor; preCFU-E, pre-erythrocyte colony-forming units; preGM, pre-granulocyte/macrophage progenitors; preMegE, pre-megakaryocyte/erythrocyte progenitors; CLP, common lymphoid progenitor.

**Figure S8.**
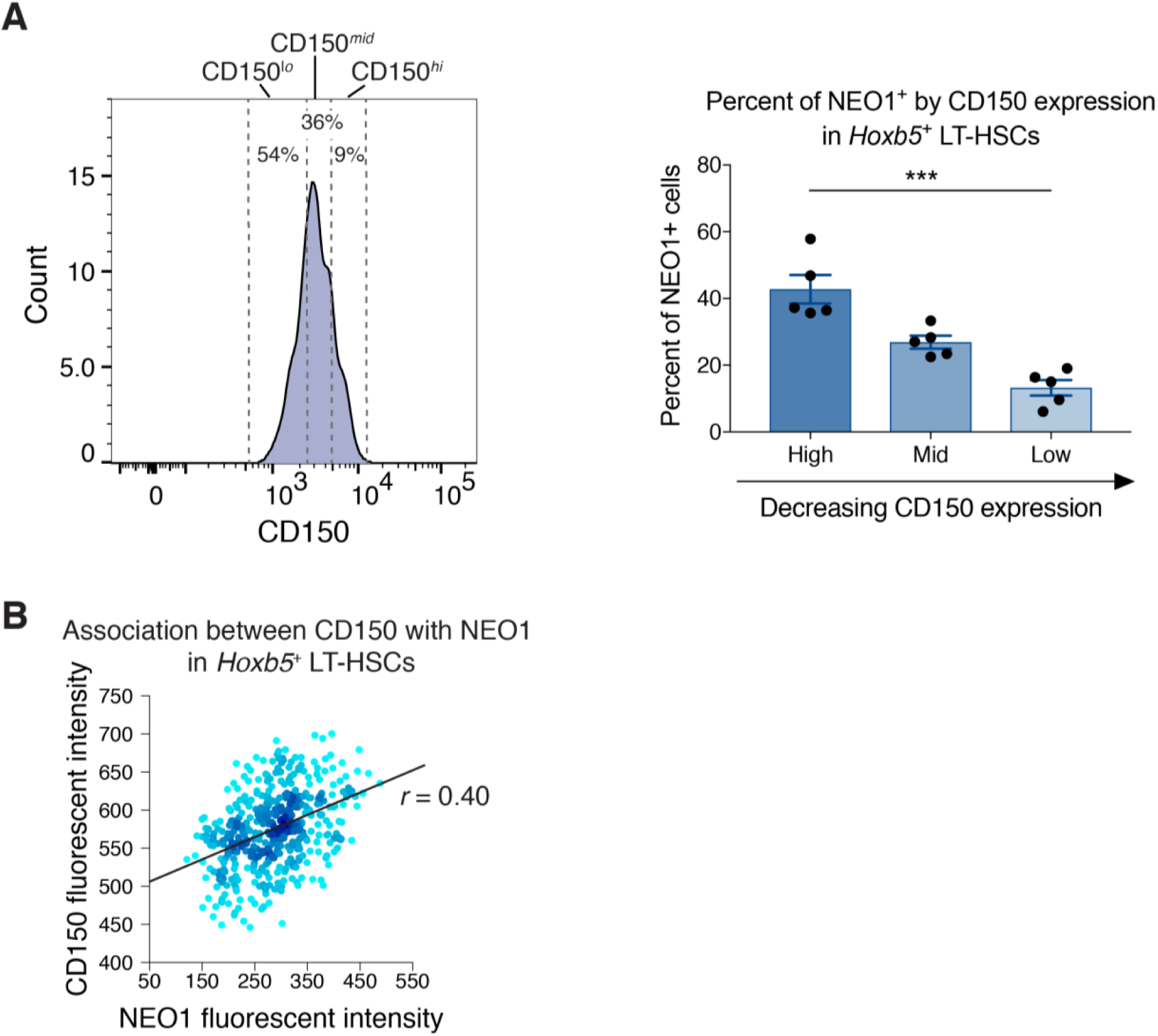
Association between NEO1 and CD150 in *Hoxb5*^+^ LT-HSCs by flow cytometry. (**A**) Histogram (*left*) and bar plots (*right*) showing the percent of NEO1^+^ cells in three bins of CD150 expression, including ‘High’, ‘Mid’, and ‘Low’. Statistical significance was calculated by a paired, two-tailed Student’s *t*-test between ‘High’ and ‘Low’. \*\*\**P* < 0.001. Bars and error bars (*right*) indicate mean ± SEM. (**B**) Representative flow cytometry diagram of NEO1 fluorescent intensity (*x*-axis) and CD150 fluorescent intensity (*y*-axis) with a linear regression line and Pearson correlation coefficient shown.

**Figure S9.**
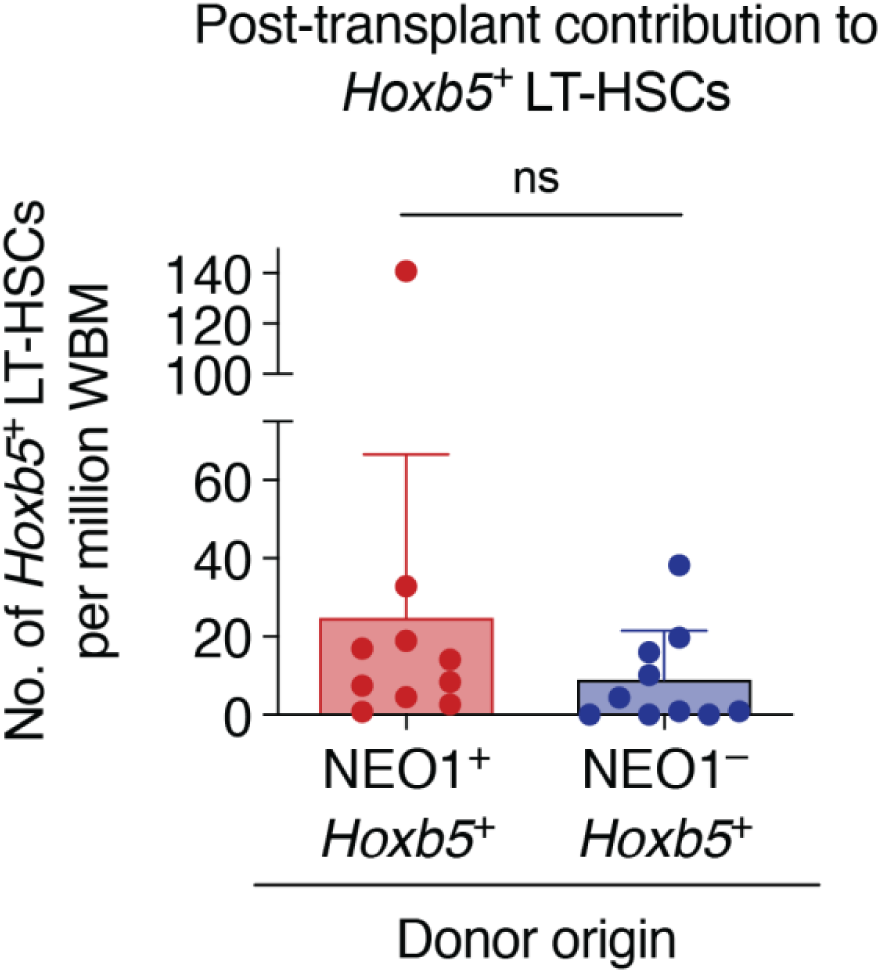
Post-transplant contribution of NEO1^+^ and NEO1^−^ *Hoxb5*^+^ LT-HSCs to *Hoxb5+ LT-HSCs*. Bar plot showing the number of *Hoxb5*^+^ LT-HSCs per million whole bone marrow (WBM) cells from each donor population 20 weeks after co-transplant of NEO1^+^ and NEO1^−^ *Hoxb5*^+^ LT-HSCs (*n* = 10). Bars and error bars indicate mean ± SD. Statistical significance was calculated by a paired, two-tailed Student’s *t*-test. ns = not significant, *P* ≥ 0.05.

